# Simple and efficient measurement of transcription initiation and transcript levels with STRIPE-seq

**DOI:** 10.1101/2020.01.16.905182

**Authors:** Robert A. Policastro, R. Taylor Raborn, Volker P. Brendel, Gabriel E. Zentner

## Abstract

Accurate mapping of transcription start sites (TSSs) is key for understanding transcriptional regulation. However, current protocols for genome-wide TSS profiling are laborious and/or expensive. We present Survey of TRanscription Initiation at Promoter Elements with high-throughput sequencing (STRIPE-seq), a simple, rapid, and cost-effective protocol for sequencing capped RNA 5’ ends from as little as 50 ng total RNA. Including depletion of uncapped RNA and SPRI bead cleanups, a STRIPE-seq library can be constructed in about five hours. We demonstrate application of STRIPE-seq to TSS profiling in yeast and human cells and show that it can also be effectively used for measuring transcript levels and differential gene expression analysis. In conjunction with our ready-to-use computational analysis workflows, STRIPE-seq is a straightforward, efficient means by which to probe the landscape of transcriptional initiation.

## Introduction

Transcription initiation is a critical control point for gene expression, integrating numerous regulatory inputs that converge on promoters and enhancers to fine-tune transcriptional activity. Understanding the spatiotemporal control of transcriptional initiation hinges upon accurate identification of transcription start sites (TSSs) and co-regulated clusters of TSSs, commonly referred to as transcription start regions (TSRs). Usage of alternative TSSs is widespread (Davuluri et al., 2008; Reyes and Huber, 2018) and typically results in mRNAs with shortened or lengthened 5’ untranslated regions (5’ UTRs). Alternative TSS selection can lead to the inclusion or exclusion of upstream open reading frames (uORFs), which alter mRNA stability and translational efficiency (Barbosa et al., 2013; Calvo et al., 2009; Kurihara et al., 2018; Wang et al., 2016). Differential TSS usage is also known to give rise to alternative protein products from the associated downstream gene. For instance, use of alternative TSSs for the maize *mybr35* gene in distinct tissues controls the inclusion or exclusion of a sequence encoding a zinc finger in the associated transcript (Mejía-Guerra et al., 2015). An example of profound phenotypic differences that can underlie alternative promoter usage occurs at the mouse *Agouti* locus, wherein usage of alternative promoters produces transcripts with distinct 5’ UTRs that confer differing temporal and regional expression, leading to the distinct pigmentation of the dorsal and ventral regions of the animal (Vrieling et al., 1994). Large-scale shifts in TSS usage are also prevalent in various developmental contexts (Adiconis et al., 2018; Batut et al., 2013; Danks et al., 2018; Haberle et al., 2014; Zhang et al., 2017) as well as human cancers (Demircioğlu et al., 2018; Thorsen et al., 2011) and inflammatory bowel diseases (Boyd et al., 2018). However, the functional relevance of these changes has been only minimally explored.

Due to the importance of TSS selection to the regulation of gene expression, several methods for global TSS profiling have been developed. The most frequently cited method is Cap Analysis of Gene Expression (CAGE) (Shiraki et al., 2003), wherein total RNA is reverse transcribed and 5’-complete cDNA:RNA hybrids are isolated via oxidation and biotinylation of the 5’ 7-methylguanosine (m^7^G) and streptavidin pulldown followed by the generation of adapter-ligated cDNA libraries. In conjunction with high-throughput sequencing, CAGE has been extensively used to characterize the landscape of transcription initiation across numerous species (Andersson et al., 2014; Consortium and CLST, 2014; Hoskins et al., 2011; Nepal et al., 2013; Valen et al., 2009)}(Kurihara et al., 2018; Lizio et al., 2017). Despite numerous revisions to the CAGE protocol over the years since its introduction (Kodzius et al., 2006; Murata et al., 2014; Takahashi et al., 2012), the method still remains costly and laborious, and there is still a high total RNA input requirement. The standard CAGE protocol has recently been modified to accommodate low input by adding selectively degradable carrier oligos as Super Low-Input Carrier CAGE (SLIC-CAGE), but this approach further increases the complexity, cost, and time associated with the protocol (Cvetesic et al., 2018) (see Supplemental Figure 23 and Supplemental Table 4 for cost and time estimates of TSS profiling methods).

A number of other TSS profiling methods use an oligo capping approach, which involves enzymatic removal of the 5’ m^7^G cap and replacement with a synthetic oligo, allowing selection of 5’-complete cDNAs (Suzuki and Sugano, 2003; Wakaguri et al., 2007; Yamashita et al., 2011). Methods incorporating oligo capping include Paired-End Analysis of TSSs (PEAT) (Ni et al., 2010), transcript leader sequencing (TL-seq) (Arribere and Gilbert, 2013), transcript isoform sequencing (TIF-seq) (Pelechano et al., 2013), CapSeq (Gu et al., 2012), Simultaneous Mapping Of RNA Ends (SMORE-seq) (Park et al., 2014), and Global/Precision Run-On sequencing of capped RNAs (GRO-/PRO-cap) (Core et al., 2014)}. However, oligo capping methods suffer from drawbacks including high input RNA requirements (e.g., 30 µg in the case of *Arabidopsis* PEAT (Morton et al., 2014)) and the sequence biases of RNA ligases used to attach oligo caps (Hafner et al., 2011; Jayaprakash et al., 2011). A potential barrier to entry for ligation-based methods has been the commercial discontinuation of tobacco acid pyrophosphatase (TAP), used in the original iterations of these methods for removal of the m^7^G cap. However, a number of TAP alternatives have been put forth, including the commercially available plant acid pyrophosphatase Cap-Clip (available from Cellscript, Madison, WI), the bacterial RNA 5’-pyrophosphohydrolase RppH, and a fusion of the *Schizosaccharomyces pombe* Dcp1/Dcp2 decapping complex to its activator peptide Edc1 (Paquette et al., 2018). RppH has been reported to be compatible with both the PRO-cap and Start-seq protocols (Mahat et al., 2016; Scheidegger et al., 2019).

Another staple molecular approach in TSS mapping is template-switching reverse transcription (TSRT), which leverages the propensity of MMLV-derived reverse transcriptases to act as terminal transferases, adding a few non-templated nucleotides, usually 3-4 Cs, when they reach the capped 5’-end of RNA molecules (Schmidt and Mueller, 1999). A template-switching oligo (TSO) bearing three riboguanosine residues (rGrGrG) at its 3’-end can then anneal to this CCC overhang, allowing template switching to add an adapter sequence to the 5’-end of the cDNA (Zhu et al., 2001). TSS mapping methods incorporating TSRT include 5’ serial analysis of gene expression (5’ SAGE) (Zhang and Dietrich, 2005), nano-cap analysis of gene expression (nanoCAGE 2010) (Plessy et al., 2010), single-cell tagged reverse transcription (STRT) (Islam et al., 2011), and RNA Annotation and Mapping of Promoters for the Analysis of Gene Expression (RAMPAGE) (Batut et al., 2013). TSRT provides additional specificity for mRNA 5’ ends; however, secondary template switching events may occur when reverse transcriptase reaches the end of the TSO, leading to the formation of TSO concatemers that reduce the overall complexity of the final library (Kapteyn et al., 2010; Turchinovich et al., 2014).

A newer generation of TSS profiling methods, nanoCAGE 2017 (Poulain et al., 2017), Tn5Prime (Cole et al., 2018), and Parallel Analysis of RNA 5’ Ends from low input (nanoPARE) (Schon et al., 2018), combine TSRT with simultaneous fragmentation and adapter tagging (tagmentation) by Tn5 transposase (Adey et al., 2010) to generate cDNA fragments of a size compatible with Illumina sequencing. However, commercially-available Tn5 is expensive, and purification of active Tn5 is laborious. nanoCAGE and nanoPARE also use custom sequencing primers, complicating pooling of other sample types in the same sequencing lane. Lastly, these methods may also suffer from the presence of TSO concatemers in the final sequencing library, although this is partially mitigated by semi-suppressive PCR in the original version of the non-Tn5-based nanoCAGE protocol (Plessy et al., 2010).

Thus, despite the successful application of TSS profiling methods to identify transcriptionally active elements on a genome-wide basis (Andersson et al., 2014; Djebali et al., 2012), their wider adoption is limited by barriers of expense, technical difficulty, and time. To overcome these hurdles, we introduce a new method, entitled Survey of TRanscription Initiation at Promoter Elements with high-throughput sequencing (STRIPE-seq). STRIPE-seq addresses several concerns of efficiency and bias inherent in other methods through a specially designed TSO, a stringent bead purification scheme, and various other methodological considerations (Figure 1; Supplemental Figure 1). Requiring only a TSRT reaction and PCR amplification following enzymatic depletion of uncapped RNA, STRIPE-seq is a simple and cost-effective protocol that can be performed in any molecular biology laboratory in approximately half a working day. In addition to the simple methodology of STRIPE-seq, we provide an end-to-end bioinformatic analysis workflow (available via GitHub code download or as a containerized package in a ready-to-use Singularity image (Kurtzer et al., 2017) to facilitate interoperability for straightforward and reproducible analysis of STRIPE-seq data. We performed STRIPE-seq in the budding yeast *Saccharomyces cerevisiae*, generating near-single base resolution maps of transcription initiation under conditions of normal growth and diamide-induced oxidative stress. In addition to profiling initiation, we find that STRIPE-seq performs comparably to standard RNA-seq in measuring transcript abundance and differential gene expression in yeast. We also demonstrate that STRIPE-seq effectively profiles TSS usage and transcript abundance in human cells. STRIPE-seq libraries can be constructed from as little as 50 ng of total RNA without modification to the base protocol, suggesting that this method can be used for profiling initiation in precious clinical or developmental samples. We envision that the simplicity of the STRIPE-seq protocol, in conjunction with the ready-to-use computational workflow, will lead to the widespread adoption of TSS profiling as a standard approach in studies of transcriptional regulation.

**Figure 1.**
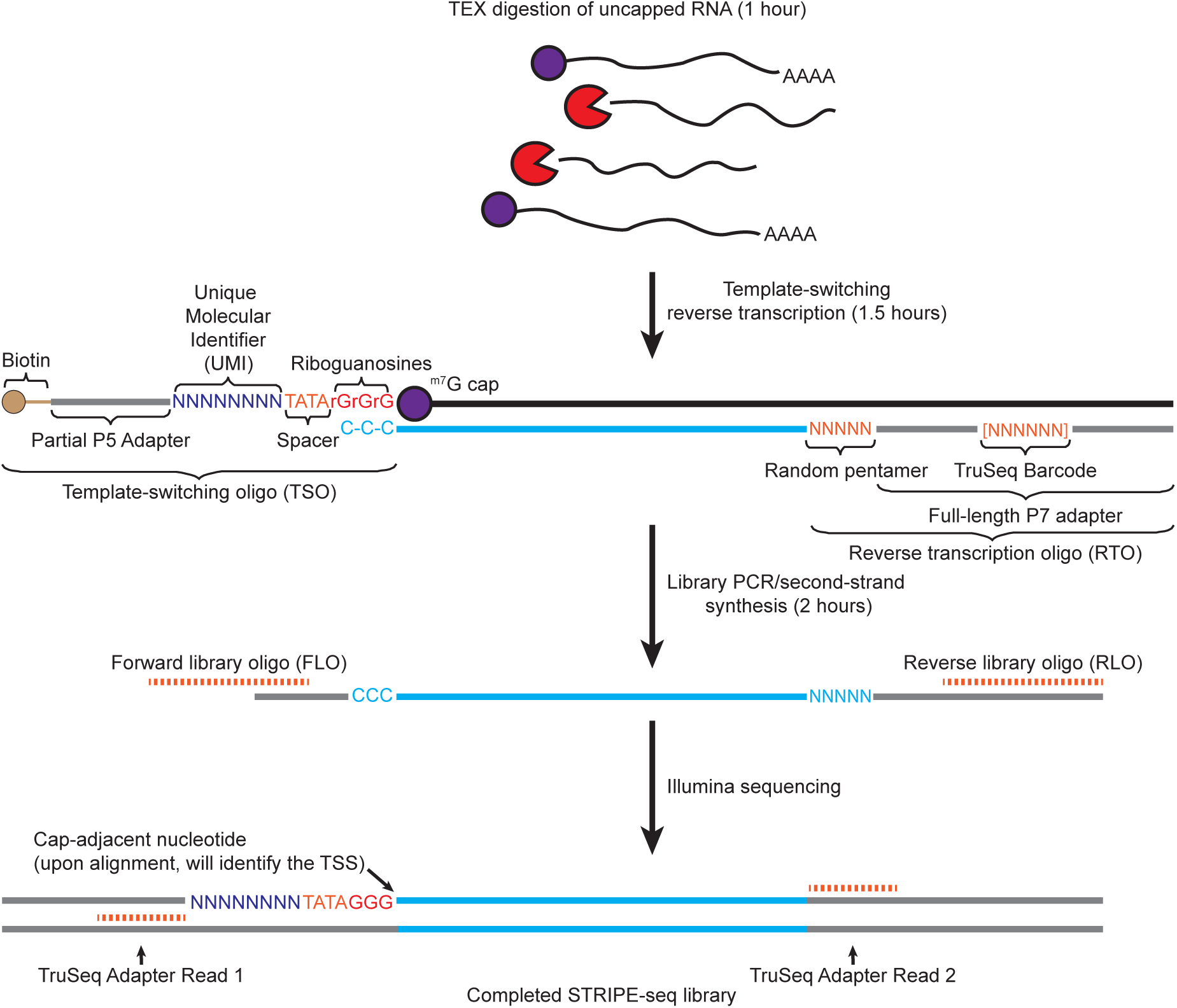
STRIPE-seq workflow. Schematic illustration of the STRIPE-seq method. Briefly, total RNA is treated with Terminator 5’-Phosphate-Dependent Exonuclease (TEX) to reduce the proportion of uncapped RNA present in the sample. After a one hour incubation, template-switching reverse transcription (TSRT) is performed using a barcoded reverse transcription oligo (RTO) primed with a random pentamer, followed by the addition of a unique molecular identifier (UMI)-containing, 5’-biotin-modified template-switching oligo (TSO) with three 3’ riboguanosines that permit the annealing of the oligo to the untemplated triplet Cs that are generated by reverse transcriptase when it reaches the m^7^G cap. Library PCR is then performed using the TSRT product as input, which ensures that TruSeq adapters are present on both sides of the insert (blue line). The cap-adjacent base, which identifies the transcription start site (TSS), is identified using the signature sequence [N_8_]-TATAGGG on the R1 read. Once the library PCR step is complete, the STRIPE-seq library is then submitted for sequencing on an Illumina platform. Timing estimates shown include bead cleanups. Please see Methods for a detailed description of the protocol.

### Design of STRIPE-seq

STRIPE-seq library construction relies on three enzymatic steps: depletion of uncapped RNA (predominantly rRNA) with Terminator 5’-phosphate-dependent exonuclease (TEX), TSRT, and library PCR. For TEX treatment, total RNA was digested in a 2 µL reaction volume, and this reaction was directly input into the TSRT reaction. For TSRT, we designed a custom reverse transcription oligonucleotide (RTO) and TSO. The RTO contains two parts: a 5-nt random sequence to facilitate random priming within transcripts followed by the full length Illumina TruSeq P7 barcode adapter (Supplemental Figure 2). The barcode was included in the RTO rather than the TSO, where it was included in RAMPAGE (Batut et al., 2013), to facilitate standard Illumina demultiplexing. Furthermore, shorter TSOs have also been shown to promote more efficient TSRT (Zajac et al., 2014).

We designed the TSO, the core of which is the Illumina TruSeq P5 Universal Adapter, with several features to address potential sources of bias (Supplemental Figure 2). First, a random 8-nt unique molecular identifier (UMI) (Kivioja et al., 2011) is positioned downstream of the P5 primer binding site. The positioning of the UMI immediately downstream of the P5 primer binding site partially addresses the low-diversity problem of Illumina sequencing, in which samples with homogenous 5’ nucleotide compositions display reduced cluster detection and subsequent loss of information (Krueger et al., 2011; Mitra et al., 2015). Furthermore, while all the data presented here were obtained with paired-end sequencing, the UMI could also be used for computational detection and removal of PCR duplicates, facilitating more accurate TSR quantification with single-end sequencing. Immediately downstream of the UMI, we placed a 4-nt spacer (TATA), which has been shown to suppress TSO invasion (*i*.*e*., annealing of the TSO to a poly-C tract in an incompletely reverse-transcribed first strand cDNA molecule) (Tang et al., 2013). We did not replace the 3’-most riboguanosine residue of the TSO with a locked nucleic acid guanylate residue as was done in Smart-seq2 because, while this substitution can increase cDNA yield (Picelli et al., 2013), it also increases strand invasion artifacts (Harbers et al., 2013). We also modified the 5’-end of the TSO with biotin to prevent the formation of large quantities of TSO concatemers that severely reduce library complexity, which occurs following secondary TS events that may occur when reverse transcriptase reaches the end of the initial TSO (Kapteyn et al., 2010; Turchinovich et al., 2014). TSO introduction into the RT reaction was withheld until 5 min into the extension step, allowing synthesis of 5’-complete first-strand cDNA to reduce TSO invasion and subsequent internal priming (Turchinovich et al., 2014). We also note that the introduction of barcodes during TSRT facilitates pooling of samples prior to library PCR.

Following TSRT, a modest SPRI (Solid Phase Reversible Immobilization) bead-based size selection is performed to remove TSO/RTO dimers and small fragment inserts. After second-strand cDNA synthesis, library PCR, and a second double-sided reaction cleanup/size selection step, the library is ready for Illumina sequencing. As STRIPE-seq relies upon bead-based size selection to optimize the size distribution of the final library, only a single round of PCR is necessary, compared with the two rounds of PCR used in tagmentation-based TSS mapping methods such as nanoCAGE, Tn5Prime, and nanoPARE.

### STRIPE-seq provides high-resolution maps of the yeast initiation landscape

We first tested the STRIPE-seq library construction protocol with three relatively modest quantities of total RNA (50, 100, and 250 ng) from the budding yeast strain S288C. Libraries constructed from RNA extracted from three independent cultures per input amount were highly consistent, with a size distribution from ∼200-1000 bp and a library amount of 25-100 ng with very little oligo dimer present (Supplemental Figure 3). No library was generated in control reactions lacking input RNA, indicating that the bio-TSO has a low potential for concatemerization and generation of artifactual libraries (Supplemental Figure 3). We then sequenced each library and performed a number of quality control steps prior to and after alignment using our specifically developed computational workflow, GoSTRIPES (Supplemental Figure 4). First, read quality was assessed and adapter sequences were trimmed. We subsequently removed reads mapping to rRNA. We observed levels of rRNA contamination ranging from 29-33% for 50 ng samples, 44.4-49.7% for 100 ng samples, and 51.8-59.3% for 250 ng samples (Supplemental Table 1). Lastly, we discarded 5’ reads without the UMI-TATAGGG sequence and trimmed it from remaining reads prior to alignment. After alignment, 92.8-95.3% of reads uniquely mapped to the genome. Aligned reads were then subjected to further quality control. First, duplicate reads, reads missing mate pairs, and non-primary alignment reads were removed. Only 13.8-55% of uniquely mapped reads qualified as likely independent, unique data points. The prevalence of PCR duplicates greatly decreased as input increased from 50 to 250 ng of total RNA, suggesting low RNA input as a limiting factor in library complexity relative to TSRT efficiency. To also assess the potential contribution of over-sequencing to the prevalence of PCR duplicates, we performed saturation analysis to computationally infer library complexity. With all methods analyzed, diminishing returns were observed above ∼1.5-2 million mappable fragments (sequenced fragments that can be mapped to the genome following computational removal of reads corresponding to rRNA), suggesting over-sequencing of STRIPE-seq libraries (Supplemental Figure 5). Furthermore, library complexity increased as RNA input increased in STRIPE-seq, further indicating that total RNA input is a limiting factor in library complexity, presumably due to the efficiency of TSRT (Wulf et al., 2019). A caveat to this analysis is that duplicate removal in SLIC-CAGE and nanoCAGE samples was not possible, making accurate determinations of library complexity difficult for those samples. After initial filtering of the alignments, we further cleaned the data by removal of read pairs with more than 3 soft-clipped bases adjacent to the presumed TSS. We allowed for the potential presence of introns in our data by accepting any read pairs containing a read with a within-read gap between 50 and 1000 nt (corresponding to a gapped alignment of a single read) as well as a maximum TLEN of 1500 (as yeast introns are infrequent and small). Only a few thousand reads per sample were removed during these final scrubbing procedure (Supplemental Table 1). Final accepted read pairs numbered between 270,815 and 739,857, which amounts to 8.8-20.4% of the initial input pairs.

To determine a read number threshold for STRIPE-seq analysis, for each STRIPE-seq sample we assessed the promoter-proximal fraction of R1 5’-ends, ideally representing the 5’-most base of the transcript and thus its TSS, with a promoter definition of −250 to +100 bp relative to an annotated start codon. At a threshold of 3 counts per TSS, the nine STRIPE-seq samples yielded promoter-proximal fractions of 0.723-0.88, with 4,314-5,075 genes having at least one unique TSS (Figure 2A, Supplemental Figure 6). Increasing the threshold beyond 3 counts slightly increased the promoter proximal fraction for each sample but resulted in a substantial loss of genes with a unique TSS. For instance, increasing the threshold to 4 increased the promoter-proximal fraction of 100 ng replicate 1 from 0.851 to 0.878 but decreased the number of genes with a unique promoter-proximal TSS from 4,710 to 4,354. We thus consider a threshold of 3 counts per TSS to be a suitable balance between removal of TSSs with 1-2 reads, which are more likely to be artifacts given the low promoter-proximal fractions associated with these thresholds, and retention of unique TSSs potentially associated with weakly expressed genes. Using this threshold, we assessed the reproducibility of STRIPE-seq TSS signal across biological replicates generated from the same amount of input RNA as well as between libraries generated using different input amounts. We normalized TSS counts using the trimmed mean of M-values (TMM) method (Robinson and Oshlack, 2010), determined Pearson correlation coefficients between samples, and hierarchically clustered the results. This analysis revealed robust concordance between replicates derived from a single input amount (Pearson’s *r* = 0.938-0.961) as well as strong correlation between datasets generated from different input amounts (Pearson’s *r* = 0.919-0.957) (Figure 2B). Given the strong correlations between STRIPE-seq replicate TSSs, we present the results of analysis of a single 100 ng sample in the main figures while showing analysis of all replicates in the supplement.

**Figure 2.**
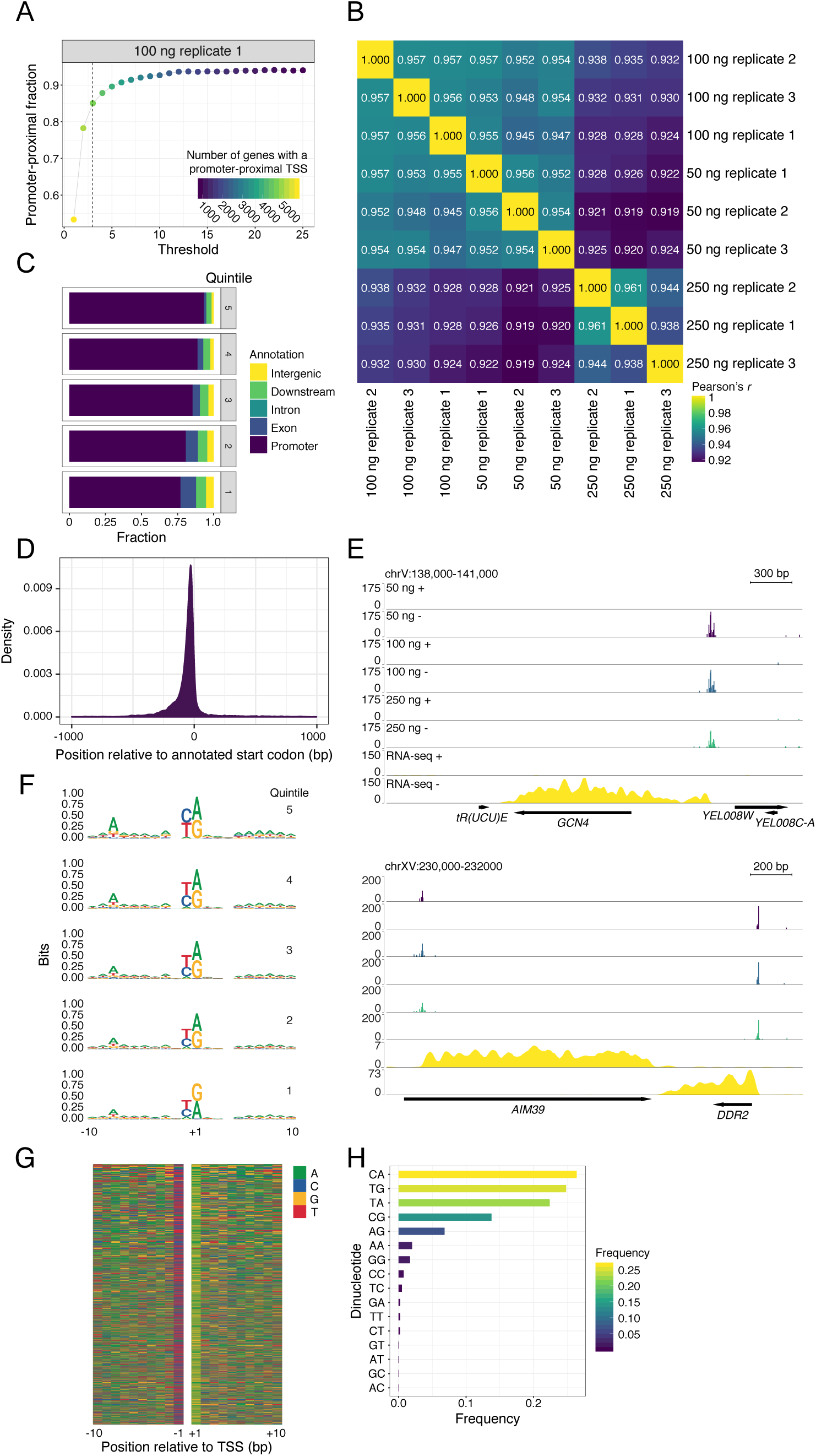
STRIPE-seq captures the yeast initiation landscape. (A) Plot of the fraction of unique TSSs that are promoter-proximal (−250 to +100 bp relative to an annotated start codon) at or above the indicated read threshold. Dot color is indicative of the number of genes with a promoter-proximal TSS. Corresponding plots for all STRIPE-seq samples are presented in Supplemental Figure 6. (B) Hierarchically clustered heatmap of Pearson’s *r* values for pairwise comparisons between TSSs identified in STRIPE-seq samples. Prior to clustering, samples were thresholded such that each TSS had to have at least 3 raw counts in one sample and then TMM normalized. (C) Genomic distribution of TSSs in 100 ng STRIPE-seq replicate 1 broken into quintiles by TSS strength. Genomic distributions of TSSs for all STRIPE-seq samples are presented in Supplemental Figure 7. (D) Density plot of 100 ng STRIPE-seq replicate 1 TSSs relative to annotated start codons. (E) Genome browser tracks showing CPM-normalized STRIPE-seq (replicate 1 for each input amount) and poly(A)+ RNA-seq (replicate 1) at two representative regions of the yeast genome. (F) Sequence logos of TSSs detected in 100 ng STRIPE-seq replicate 1 broken into quintiles by TSS strength. Sequence logos of TSSs in all STRIPE-seq samples are presented in Supplemental Figure 9. (G) Nucleotide color plot of the sequence context of TSSs detected in 100 ng STRIPE-seq replicate 1. TSSs are ranked descending by read count. (H) Dinucleotide frequencies at TSSs detected in 100 ng STRIPE-seq replicate 1. Dinucleotide frequencies at TSSs in all STRIPE-seq samples are presented in Supplemental Figure 10.

We first performed a more detailed analysis of the genomic distribution of TSSs detected by STRIPE-seq. Division of TSS distribution into quintiles based on read count revealed that the strongest TSSs were most likely to be promoter-proximal, with progressively smaller promoter-proximal fractions as TSS strength decreased (Figure 2C, Supplemental Figure 7). Analysis of TSS density relative to annotated start codons also revealed a strong promoter-proximal preference (Figure 2D). We also inspected TSS signal in relation to annotated open reading frames (ORFs) and poly(A)+ RNA-seq signal at individual genomic regions. At *GCN4*, encoding a transcription factor that activates amino acid biosynthesis genes during amino acid starvation (Hinnebusch, 1997), we observed RNA-seq signal extending nearly 600 bp upstream of the start codon, where a cluster of TSSs (a transcription start region, TSR) was found (Figure 2E). This observation is consistent with translational regulation of *GCN4* mRNA by four upstream ORFs (uORFs) (Mueller and Hinnebusch, 1986). At the *AIM39* locus, we found a TSR downstream of the annotated start codon, with no upstream RNA-seq signal (Figure 2E), suggesting misannotation of the *AIM39* start codon. Indeed, comparison of STRIPE-seq and RNA-seq signal to ribosome profiling (Ribo-seq) data (Nissley et al., 2016) revealed strong ribosome occupancy downstream of the *AIM39* TSR and 5’ end of the associated RNA-seq signal (Supplemental Figure 8).

We next analyzed the sequence context of STRIPE-seq-detected TSSs. Consistent with previous work (Zhang and Dietrich, 2005), we detected a consensus A_−8_Y_−1_R_+1_ motif (Figure 2F-G, Supplemental Figure 9). Through sequence analysis of TSS quintiles divided by TSS strength, we observed that the information content of the A_−8_ base decreased as TSS strength diminished (Figure 2F), consistent with the previously described positive relationship between this position and TSS usage (Zhang and Dietrich, 2005). Lastly, we analyzed the dinucleotide frequencies at TSSs identified by STRIPE-seq. All replicates identified the four possible Y_−1_R_+1_ combinations (CA, TG, TA, and CG) as the most prevalent initiator dinucleotides (Figure 2H, Supplemental Figure 10), consistent with previous data (Lu and Lin, 2019; Malabat et al., 2015; Zhang and Dietrich, 2005). It has been previously reported in traditional CAGE and TSRT-based protocols such as nanoCAGE that extra spurious bases, especially guanosines, are sometimes present in the 5’ most base of the R1 read (which corresponds to a cytosine on the first-strand cDNA) (Cumbie et al., 2015; Cvetesic et al., 2018). Furthermore, a recent in-depth analysis of TSRT showed an almost universal addition of an extra cytosine on the cDNA adjacent to the TSS of capped RNA, but almost no addition of this base in uncapped RNA, which is speculated to occur due to the cap acting as a template for reverse transcriptase (Wulf et al., 2019). In CAGE and nanoCAGE based analysis, rates of addition for these extra bases are often inferred from those that were not incidentally templated onto the reference genome, but rather soft-clipped by the alignment software (Haberle et al., 2015). For the STRIPE-seq read pairs surviving the initial quality control steps, roughly 50-70% had soft-clipped bases on the 5’ end of the R1 read, a majority of which only had one or two additional bases added (Supplemental Figure 11). Most of the added bases on the first-strand cDNA were cytosine, with a much smaller fraction of thymidine (Supplemental Figure 12). These values are consistent with what is seen in the SLIC-CAGE and nanoCAGE samples we also analyzed for soft-clipped bases, and are consistent with previous literature. Taken together, these observations indicate that STRIPE-seq effectively and comprehensively profiles the yeast initiation landscape.

### Quantification of transcript levels with STRxIPE-seq

As the primary focus of STRIPE-seq is profiling of initiation events, our analyses thus far have focused on the 5’-most base of sequenced R1 reads. However, TSRT based methods are routinely used for general quantification of annotated transcripts in bulk and single-cell RNA-seq (Picelli et al., 2013; Turchinovich et al., 2014). We sequenced STRIPE-seq libraries in paired-end mode to facilitate simple duplicate removal and enhanced assignment of TSSs to transcripts, as each sample retains additional information on transcript origin due to the presence of reverse (i.e. R2) reads usually originating from within transcript bodies. Thus, assignment of paired-end STRIPE-seq fragments to transcripts could in principle be used to more accurately measure transcript abundance. To test this possibility, we assessed the correlation between STRIPE-seq and poly(A)+ RNA-seq signal within annotated transcripts. We observed robust correlations between all STRIPE-seq and RNA-seq samples (Spearman’s *⍴* = 0.786-0.846) (Figure 3). Correlations improved when more total RNA was used as input, likely due to capture of more moderately expressed transcripts.

**Figure 3.**
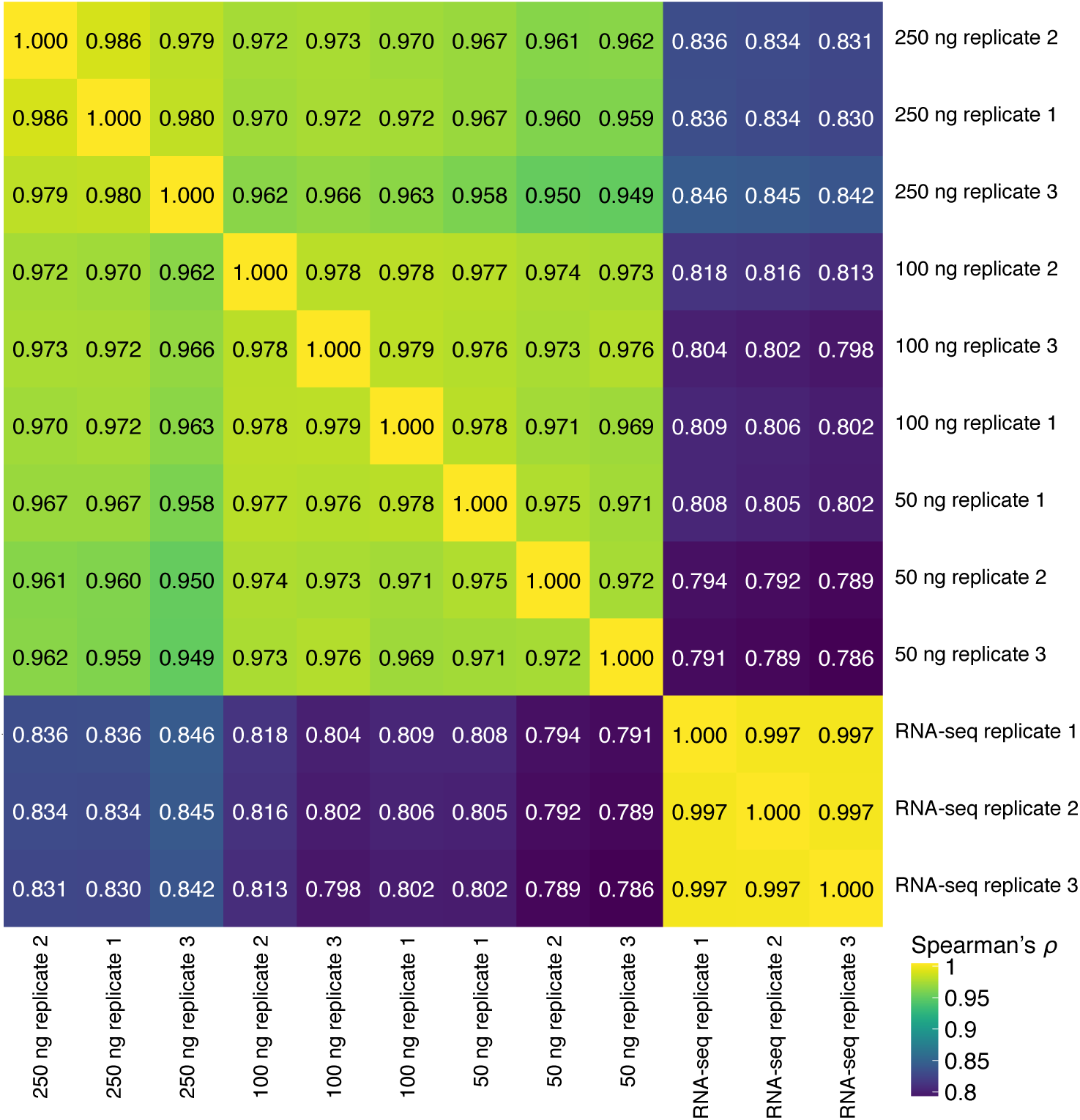
STRIPE-seq provides RNA-seq-like information on transcript abundance. Hierarchically clustered heatmap of Spearman’s *⍴* values for pairwise comparisons between TMM-normalized per-gene STRIPE-seq and poly(A)+ RNA-seq fragment counts.

### Systematic comparison of yeast STRIPE-seq with CAGE-based methods

A previous study used three distinct iterations of CAGE (nanoCAGE, nAnT-iCAGE, and SLIC-CAGE) (Cvetesic et al., 2018) to profile the initiation landscape of yeast. This work established high concordance between reference nAnT-iCAGE data and SLIC-CAGE while suggesting inferior performance of nanoCAGE in detecting TSSs. We thus set out to systematically compare STRIPE-seq to these methods. To this end, we compared our nine STRIPE-seq samples to two replicate 100 ng SLIC-CAGE datasets, two replicate datasets for nanoCAGE using 500 ng total RNA with TEX treatment, and two replicates of nanoCAGE using 25 ng input RNA without TEX treatment. As with STRIPE-seq, a threshold of at least 3 counts per TSS provided a suitable balance between promoter-proximal TSS fraction and number of genes with a unique TSS in SLIC-CAGE and nanoCAGE datasets (Supplemental Figure 13). Duplicate removal was not possible for SLIC-CAGE or nanoCAGE, as both methods were sequenced in single-end mode and the UMI for the nanoCAGE samples was removed prior to deposition. We first assessed correlation between all three methods in a conservative promoter window of −250 to +100 bp relative to the annotated start codons of 6,572 mRNA transcripts. We observed good correlation between STRIPE-seq and SLIC-CAGE signal at promoters (Spearman’s *⍴* = 0.708-0.743) (Figure 4A). Interestingly, though STRIPE-seq and nanoCAGE both rely on TSRT, correlations between STRIPE-seq and nanoCAGE were weaker than those between STRIPE-seq and SLIC-CAGE (Spearman’s *⍴* = 0.609-0.680) (Figure 4A). Consistent with these results, visual inspection of TSS signal at selected loci revealed high similarity between STRIPE-seq and SLIC-CAGE, with similar patterns of TSS distributions and higher TSR complexity relative to nanoCAGE (Figure 4B).

**Figure 4.**
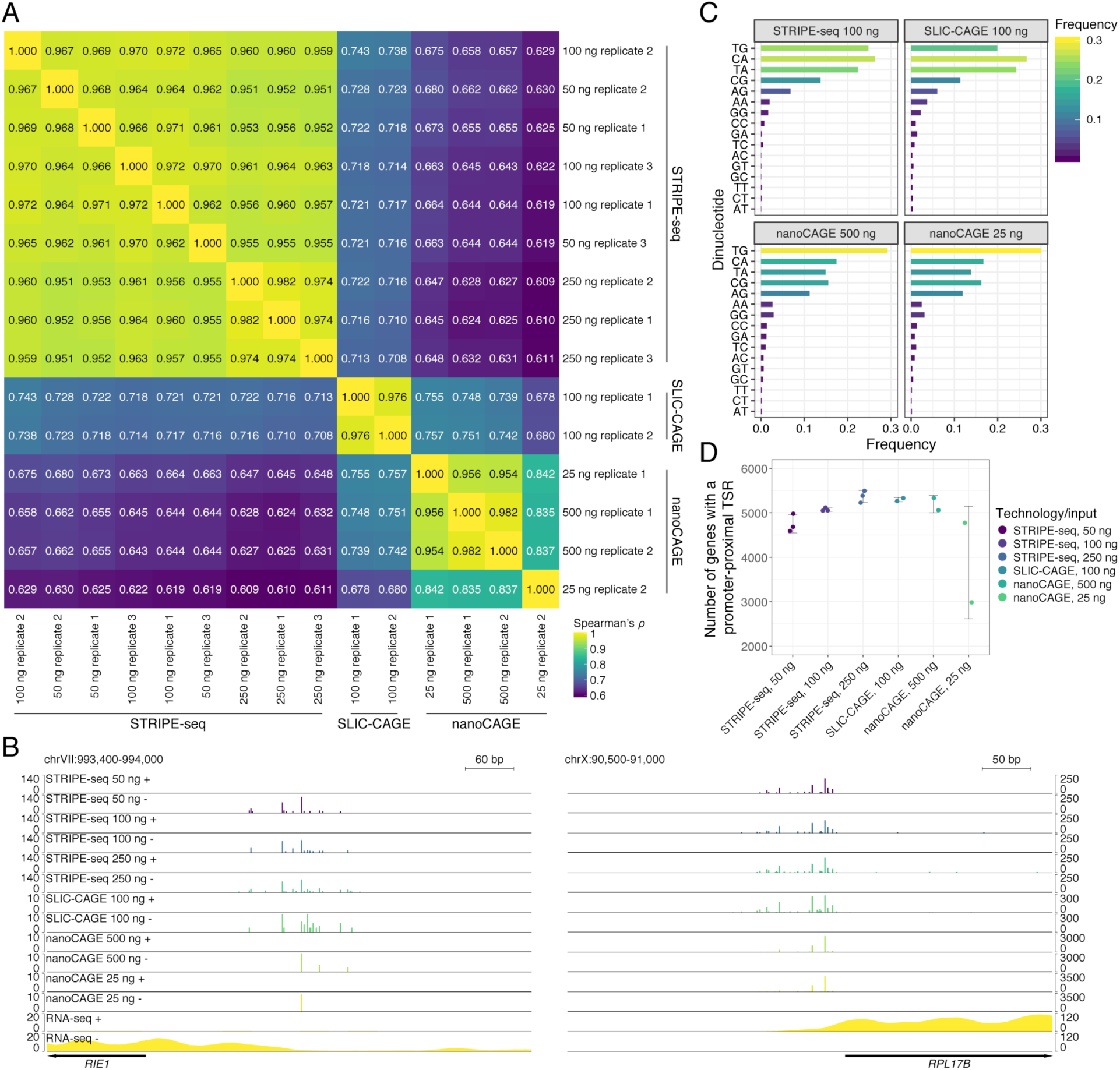
Comparison of yeast STRIPE-seq to SLIC-CAGE and nanoCAGE. (A) Hierarchically clustered heatmap of Spearman’s *⍴* values for pairwise comparisons of STRIPE-seq, SLIC-CAGE, and nanoCAGE signal within promoter regions (−250 to +100 bp relative to an annotated start codon). Prior to clustering, samples were thresholded such that each promoter had to have at least raw 3 counts in one sample, and then counts were TMM normalized. (B) Genome browser-style tracks showing CPM-normalized STRIPE-seq, SLIC-CAGE, nanoCAGE (replicate 1 for each input amount) and poly(A)+ RNA-seq (replicate 1) at two representative regions of the yeast genome. (C) Dinucleotide frequencies at TSSs detected in the indicated samples. Dinucleotide frequencies at TSSs in all replicates of all technologies are presented in Supplemental Figure 14. (D) Jitter plot of the number of genes with a promoter-proximal TSR in each sample. Error bars represent the standard deviation.

We next compared the dinucleotide frequencies of TSSs detected by STRIPE-seq, SLIC-CAGE, and nanoCAGE. STRIPE-seq and SLIC-CAGE both preferentially detected all four Y_−1_R_+1_ combinations (CA, TG, TA, and CG), with CA most preferred (Figure 4C, Supplemental Figure 14). STRIPE-seq TSSs displayed a slight preference for TG versus SLIC-CAGE TSSs, which were slightly more likely to have a TA dinucleotide. The overall dinucleotide frequency distribution of STRIPE-seq TSSs was much more similar to that of SLIC-CAGE TSSs versus nanoCAGE TSSs, which displayed a strong bias for a TG dinucleotide (Figure 4C, Supplemental Figure 14). We next assessed the numbers of genes with a detectable TSR in each method as a means to analyze methodological sensitivity. We associated TSRs with transcripts and then counted the number of transcripts with a promoter-proximal TSR in each sample. At 50, 100, and 250 ng of total RNA input, STRIPE-seq detected 4,591-4,980, 5,049-5,119, and 5,228-5,494 such transcripts, respectively. With 100 ng input RNA, SLIC-CAGE detected 5,265 and 5,330 transcripts with promoter-proximal TSRs, while nanoCAGE detected 5,057 and 5,334 transcripts with a promoter-proximal TSRs at 500 ng and 2,985 and 4,775 such transcripts at 25 ng (Figure 4D). We conclude that the sensitivity of STRIPE-seq in TSR discovery is comparable to that of SLIC-CAGE and higher-input nanoCAGE in yeast. We note that a potential limitation of these comparisons between STRIPE-seq, SLIC-CAGE, and nanoCAGE is biological variability: we performed STRIPE-seq in the S288C strain, while both previously published CAGE datasets used RNA derived from strain BY4741, an auxotrophic derivative of S288C.

### Differential STRIPE-seq analysis identifies changes in yeast TSR usage and transcript abundance

Thus far, we have shown that STRIPE-seq robustly detects TSSs under normal growth conditions. As a major potential application of this method is the detection of changes in TSS usage between distinct biological conditions, we investigated the capability of STRIPE-seq to detect TSRs altered by oxidative stress. We therefore treated yeast with the thiol-reactive chemical diamide, extracted total RNA, and performed STRIPE-seq. Correlation analysis of TSSs detected in three untreated 100 ng STRIPE-seq replicates and three 100 ng STRIPE-seq replicates derived from cells treated with 1.5 mM diamide for 1 hr revealed robust within-group concordance (untreated Pearson’s *r* = 0.957-0.958, diamide Pearson’s *r* = 0.942-0.963), but greatly reduced correlation between conditions (Pearson’s *r* = 0.635-0.654) (Figure 5A), suggesting a widespread shift in TSS usage upon diamide stress. Consistent with this, correlation of untreated and diamide-treated samples within a merged set of 4,866 TSRs was also weak compared to within-sample concordance (Pearson’s *r* = 0.651-0.679; untreated Pearson’s *r* = 0.983-0.984, diamide Pearson’s *r* = 0.963-0.989) (Figure 5A). Visualization of STRIPE-seq and matched RNA-seq signal at *HSP150*, encoding a cell wall mannoprotein upregulated by various stressors (Russo et al., 1993), revealed increased TSR usage and transcript abundance as measured by matched poly(A)+ RNA-seq (Figure 5B). At the same region we also observed downregulation of the convergent *CIS3* locus (Figure 5B). We also observed reduced TSR usage and RNA-seq signal at the convergent *RPS16B* and *RPL13A* genes (Figure 5B), consistent with the known repression of ribosomal protein gene expression during the environmental stress response (Gasch et al., 2000; Weiner et al., 2012).

**Figure 5.**
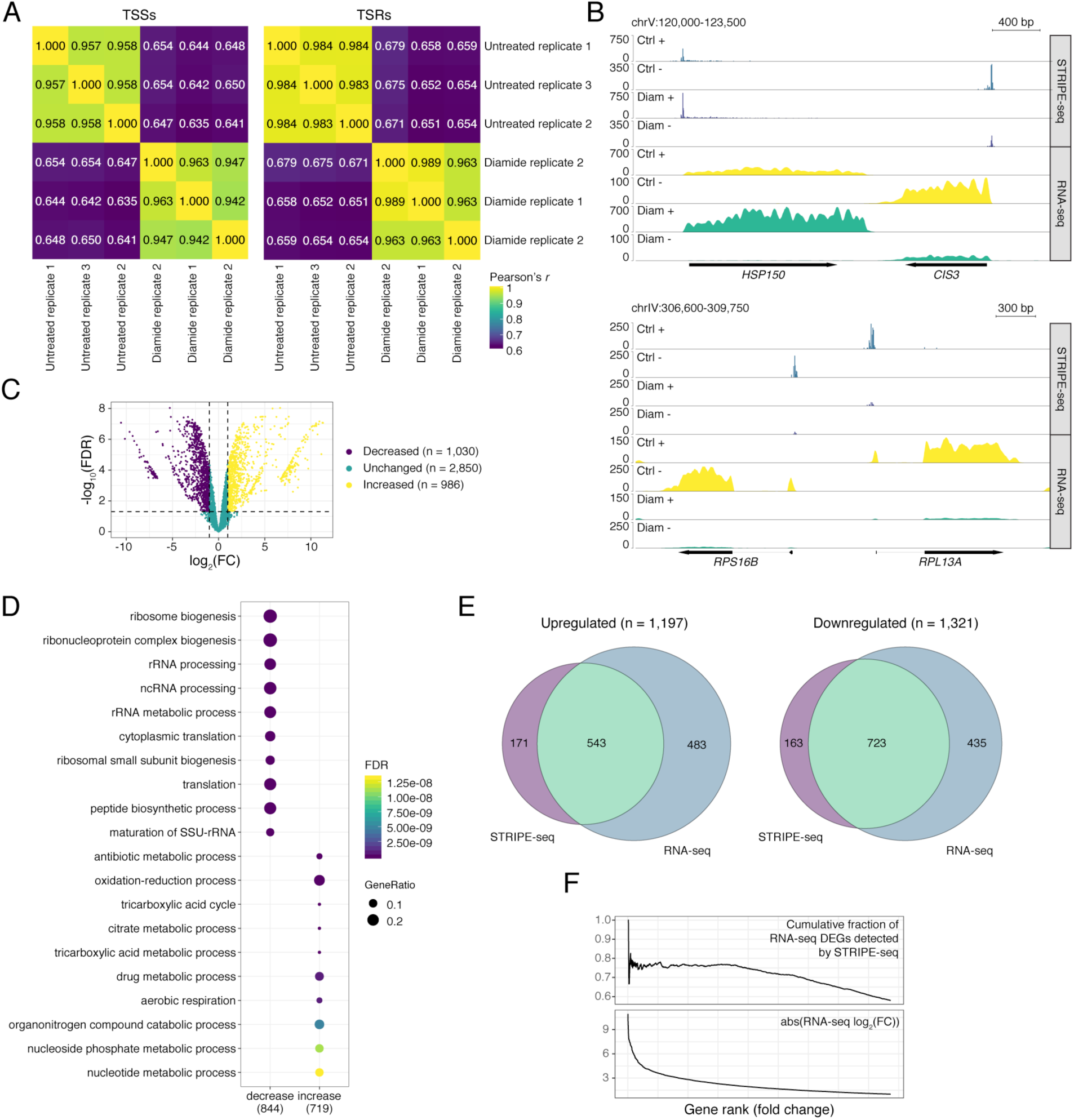
STRIPE-seq captures differential TSR usage and transcript abundance. (A) Hierarchically clustered heatmaps of Pearson’s *r* values for pairwise comparisons between merged TSS and TSR sets from 100 ng control and diamide STRIPE-seq samples. (B) Genome browser-style tracks showing CPM-normalized STRIPE-seq and poly(A)+ RNA-seq from control and diamide-treated samples at two representative regions of the yeast genome. (C) Volcano plot of differential TSRs resulting from comparison of control and diamide-treated samples. (D) Dot plots of GO biological process terms for genes associated with TSRs that increased and decreased upon diamide treatment. (E) Venn diagrams of the overlap between DEGs identified by STRIPE-seq and RNA-seq in control and diamide-treated samples. (F) Cumulative distribution plots for fractions of DEGs captured by STRIPE-seq versus log_2_(FC) in poly(A)+ RNA-seq.

To more systematically characterize differential initiation in diamide-treated yeast, we performed differential TSR analysis using the merged untreated/diamide TSR set. This analysis revealed 986 upregulated and 1,030 downregulated TSRs at a fold change cutoff of 2 and an FDR threshold of 0.05 (Figure 5C, Supplemental Table 2). Of these, 756/986 (76.7%) upregulated and 871/1,030 (84.6%) downregulated TSRs were within the −250 to +100 promoter window (Supplemental Table 2). To determine the relevance of these differential TSRs, we performed gene ontology analysis of the genes associated with promoter-proximal differential TSRs. Downregulated TSRs were strongly enriched for biological processes related to ribosome and rRNA biogenesis (Figure 5D), again consistent with the general downregulation of ribosomal protein genes observed during environmental stress (Gasch et al., 2000; Weiner et al., 2012). Consistent with the cells responding to oxidative stress, upregulated TSRs were enriched for a number of metabolic processes as well as the Gene Ontology term “oxidation-reduction process” (Figure 5D).

As we showed that quantification of STRIPE-seq fragments within transcripts provides measurements of transcript abundance comparable to those determined by RNA-seq (Figure 3), we next asked if RNA-seq-like analysis of control and diamide STRIPE-seq datasets could provide comparable results to a more conventional differential expression analysis. Correlation analysis of three 100 ng control and three 100 ng diamide-treated STRIPE-seq analysis alongside matched RNA-seq replicates again revealed robust concordance (untreated Spearman’s *⍴* = 0.798-0.818, diamide Spearman’s *⍴* = 0.842-0.850) (Supplemental Figure 15). Differential expression analysis with STRIPE-seq yielded 714 upregulated and 886 downregulated genes, while RNA-seq detected 1,026 upregulated and 1,158 downregulated genes (Supplemental Table 3). A total of 1,197 upregulated genes were detected by STRIPE-seq and/or RNA-seq; of these, 543 (45.4%) were shared, 171 (14.3%) were specific to STRIPE-seq, and 483 (40.4%) were specific to RNA-seq (Figure 5E). We detected a total of 1,321 downregulated genes, with 723 (54.7%) shared between STRIPE-seq and RNA-seq, 163 (12.3%) specific to STRIPE-seq, and 435 (32.9%) specific to RNA-seq (FIgure 5E). Despite the smaller number of differentially expressed genes (DEGs) detected by STRIPE-seq versus RNA-seq, similar biological processes were enriched in up- and downregulated genes (Supplemental Figure 16), indicating that RNA-seq-like analysis of STRIPE-seq data can accurately capture overall changes in the cellular transcriptional program. To probe whether the reduced number of DEGs reported by STRIPE-seq might be attributable to reduced sensitivity, we assessed the cumulative fraction of RNA-seq DEGs detected by STRIPE-seq as a function of the absolute value of the RNA-seq log_2_(FC). We found that STRIPE-seq captured a large fraction of genes with robust fold changes in RNA-seq but was less likely to detect DEGs with moderate to low fold changes in RNA-seq (Figure 5F).

### STRIPE-seq robustly profiles initiation and transcript abundances in human cells

To explore the utility of STRIPE-seq in analyzing more complex initiation landscapes, we performed STRIPE-seq in K562 erythroleukemia cells. We constructed three biological replicate STRIPE-seq libraries using 100 ng of total RNA. As for yeast, robust libraries with a size distribution of ∼200-1000 nt and little to no oligo dimers were generated (Supplemental Figure 3). Processing of reads with GoSTRIPES (see Methods) revealed variable levels of rRNA reads ranging from 20.5-22.7% (Supplemental Table 1). We obtained proportions of uniquely mapped reads from 89.3-92.8%, though only 16.3-29.5% reads were considered to be unique after duplicate removal. We obtained 686,981-806,174 accepted read pairs after processing, representing 11-18.8% of the initial input. As for yeast, we assessed the extent of sequencing saturation and library complexity by downsampling mapped reads and determined the number of unique genes with a promoter-proximal TSS. Our findings suggest under-sequencing of K562 STRIPE-seq libraries and that library complexity would benefit from a sequencing depth of 20-30 million mappable fragments (Supplemental Figure 17). Although not tested directly for human samples, library complexity would likely also benefit from increased input amounts as was seen in the yeast samples. For comparison, we also analyzed CAGE, RAMPAGE, and nanoCAGE-XL experiments performed in K562 cells (Adiconis et al., 2018)). As expected, CAGE was the most sensitive method, detecting upwards of 15,000 genes with a promoter-proximal TSS at < 10 million mappable fragments (Supplemental Figure 17). However, the comparison between CAGE and STRIPE-seq is not straightforward, as these CAGE libraries were constructed with the no-amplification nAnT-iCAGE approach and very high input (10 µg), while STRIPE-seq uses PCR and low input (100 ng). The most appropriate comparison for STRIPE-seq is RAMPAGE, which uses TSRT, PCR, and the cap-trapping approach used in CAGE, but with a high (5 µg) amount of RNA input. One deeply sequenced RAMPAGE sample detected a few thousand more genes with a promoter-proximal TSS than STRIPE-seq, but a second RAMPAGE replicate appeared essentially identical to STRIPE-seq in this regard (Supplemental Figure 17).

At a threshold of 3 counts per TSS, we obtained promoter-proximal fractions (defined for these samples as −500 to +500 relative to an annotated TSS) of 0.938-0.942, with 7,353-7,778 genes having at least one unique STRIPE-seq TSS (Figure 6A). K562 STRIPE-seq TSSs at a threshold of 3 were highly reproducible (Pearson’s *r* = 0.932-0.937) (Figure 6B). Consistent with the threshold analysis, the majority of detected TSSs were found within promoter regions, with stronger TSSs displaying a greater promoter bias (Figure 6C, Supplemental Figure 18A), an observation also confirmed by analysis of TSS density relative to annotated TSSs (Figure 6D). Visual inspection revealed capture of features reflective of the complexity of the human transcriptome. For instance, at the *TBPL1* locus, encoding the TATA-binding protein (TBP) paralog TBP-related factor 2 (TRF2) (Akhtar and Veenstra, 2011), we observed preferential usage of a TSR that would lead to a short form of its 5’ UTR, a finding supported by visualization of matched poly(A)+ RNA-seq data (Figure 6E). We also detected robust initiation at an internal site within the *MALAT1* gene, encoding a multifunctional long noncoding RNA (Amodio et al., 2018) (Figure 6E).

**Figure 6.**
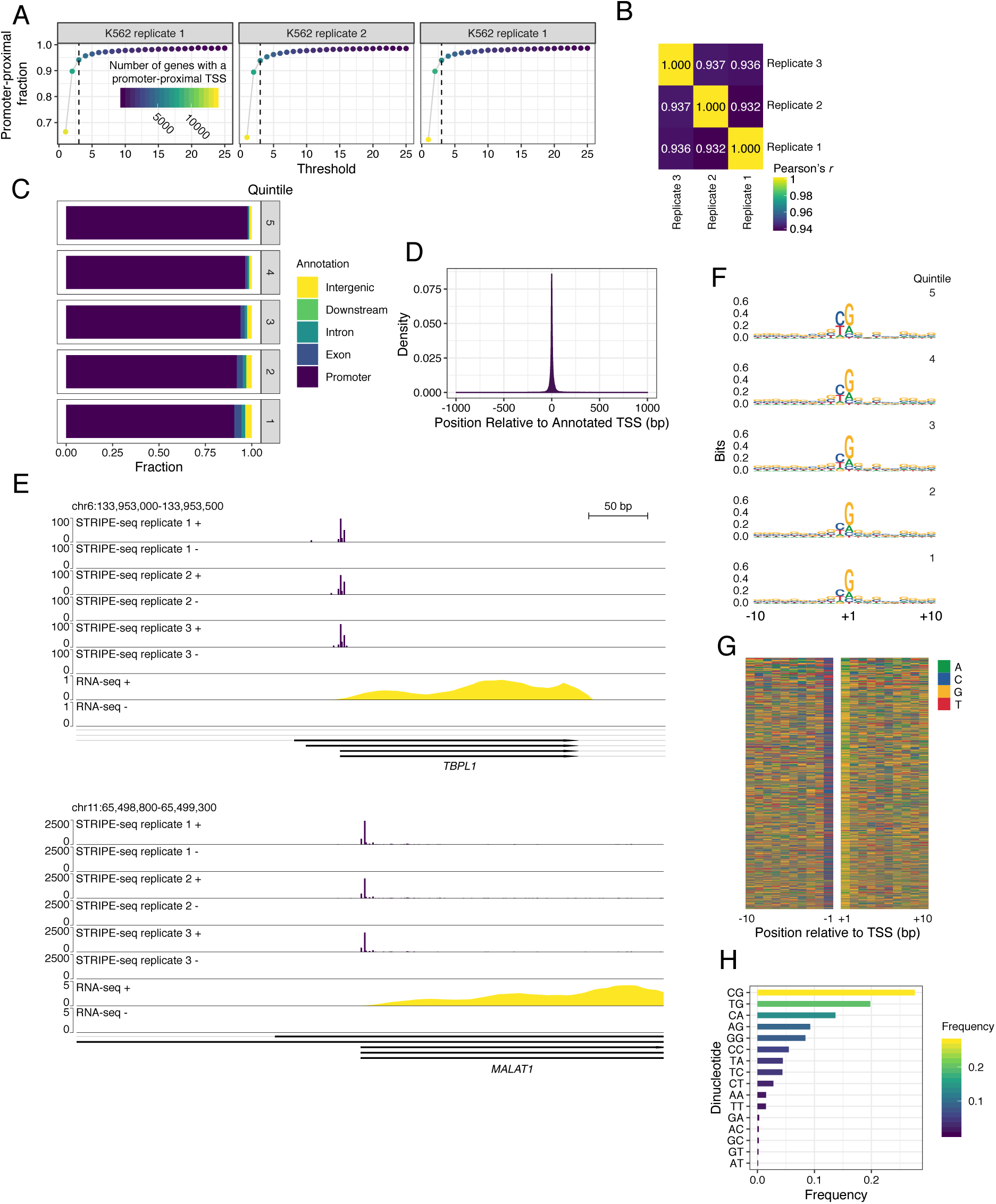
STRIPE-seq profiling of the human initiation landscape. (A) Plot of the fraction of unique TSSs that are promoter-proximal at the indicated read threshold. Dot color and size are indicative of the number of genes with a promoter-proximal TSS. (B) Heatmap of Pearson’s *r* values for pairwise comparisons between TSSs identified in STRIPE-seq samples. Prior to clustering, samples were thresholded such that each TSS had to have at least 3 raw counts in one sample, and then counts were TMM normalized. (C) Genomic distribution of TSSs in K562 STRIPE-seq replicate 1 broken into quintiles by TSS strength. Genomic distributions of TSSs for all K562 STRIPE-seq samples are presented in Supplemental Figure 18A. (D) Density plot of K562 STRIPE-seq replicate 1 TSSs relative to annotated TSSs. (E) Genome browser tracks showing CPM-normalized STRIPE-seq and poly(A)+ RNA-seq (replicate 1) at two representative regions of the human genome. (F) Sequence logos of TSSs detected in K562 STRIPE-seq replicate 1 broken into quintiles by TSS read count. Sequence logos of TSSs in all K562 STRIPE-seq samples are presented in Supplemental Figure 18B. (G) Nucleotide color plot of the sequence context of TSSs detected in K562 STRIPE-seq replicate 1. TSSs are ranked descending by read count. (H) Dinucleotide frequencies at TSSs detected in K562 STRIPE-seq replicate 1. Dinucleotide frequencies at TSSs in all STRIPE-seq samples are presented in Supplemental Figure 18C.

We then analyzed the sequences found at STRIPE-seq TSSs. Consistent with previously published CAGE data, we detected a strong Y_−1_R_+1_ (C or T and A or G) initiator with a bias for G at the +1 position (Figure 6F-G, Supplemental Figure 18B). Notably, the −1 preference for R progressively diminished as weaker TSSs were considered, while the +1 bias for R, and G in particular, remained consistent (Figure 6F). We note that while Y_−1_R_+1_ initiator we report here is consistent with that found by CAGE (Frith et al., 2008), it is distinct from that detected by 5’-GRO-seq (BBCA_+1_BW) (Vo ngoc et al., 2017), potentially due to the analysis of steady-state versus nascent transcripts by these methods, respectively. Alternatively, differences in analysis (individual TSS positions for the present analysis of STRIPE-seq data versus identification and characterization of sequences at highly focused, strong TSSs) may contribute to the different sequences detected. Consistent with sequence logo and color plot analysis, four of the five most frequently detected dinucleotides followed the NG pattern (Figure 6H, Supplemental Figure 18C). Similar to the yeast analysis, we sought to determine whether there was a strong preference for the addition of spurious guanosine(s) adjacent to putative TSSs in STRIPE-seq, CAGE, RAMPAGE, and nanoCAGE-XL. In all methods but nanoCAGE-XL, roughly 50-60% of mapped reads had soft-clipped bases, a majority of which had just a single base added (Supplemental Figure 19). Also similar to yeast, a majority of these soft clipped bases were guanosines, followed by a much smaller percentage of adenosines (Supplemental Figure 20). As human promoters tend to be more GC rich than those of yeast (Fenouil et al., 2012), it is likely more common for these spurious guanosines to be incidentally templated onto the genome, which would explain the smaller fraction of TSSs having soft-clipped bases compared to yeast in all methods.

We also assessed the ability of STRIPE-seq to measure transcript levels in K562 cells by quantification of fragments within annotated exons compared to matched poly(A)+ RNA-seq data. As observed for yeast, transcript signal for STRIPE-seq and RNA-seq was well correlated (Pearson’s *r* = 0.822-0.835) (Supplemental Figure 21), indicating that STRIPE-seq can be used for estimation of transcript abundances alongside TSS usage in human cells.

We next compared STRIPE-seq to CAGE, RAMPAGE, and nanoCAGE-XL data generated from K562 cells (Adiconis et al., 2018). We first analyzed correlation between the methods in a promoter window of −500 to +500 bp relative to 152,701 annotated TSSs of protein-coding genes. Within these promoter windows, replicates of each method were well correlated (STRIPE-seq Spearman’s *⍴* = 0.842-0.846, CAGE Spearman’s *⍴* = 0.912, RAMPAGE Spearman’s *⍴* = 0.849) (Figure 7A). STRIPE-seq signal was also highly correlated with that of CAGE (Spearman’s *⍴* = 0.790-0.805) and RAMPAGE (Spearman’s *⍴* = 0.771-0.798). Of note, these correlations are similar to that of CAGE and RAMPAGE (Spearman’s *⍴* = 0.772-0.804) (Figure 7A). Poor correlation was observed between nanoCAGE-XL and all other methods. Good correspondence between STRIPE-seq was observed visually at the shared promoter region of the *RARS2* and *ORC3* genes (Figure 7B). The dinucleotide frequencies of TSSs detected by STRIPE-seq were very similar to those found by CAGE and RAMPAGE, while no consistent dinucleotide preferences were found for nanoCAGE-XL TSSs (Figure 7C). Lastly, as was done for yeast, we assessed the sensitivity of STRIPE-seq relative to CAGE-based methods by calling TSRs and determining the number of transcripts with a promoter-proximal TSR (defined in this analysis as a TSR within −500 to +500 bp of an annotated TSS). With STRIPE-seq, we detected 8,474-9,006 transcripts with promoter-proximal TSRs, a lower quantity than were detected in the analyzed CAGE samples (13,076 and 13,903) (Figure 7D). STRIPE-seq detected fewer proximal TSR-associated transcripts than RAMPAGE replicate 1 (13,266) but more than replicate 2 (7,340), potentially due to their different sequencing depths: replicate 1 had a total of ∼36.4 M reads from two separate runs, while replicate 2 yielded ∼5.3 M reads from a single run. (Figure 7D). This observation may suggest that higher sequencing depth is necessary to fully realize the complexity of RAMPAGE libraries (Supplemental Figure 17).

**Figure 7.**
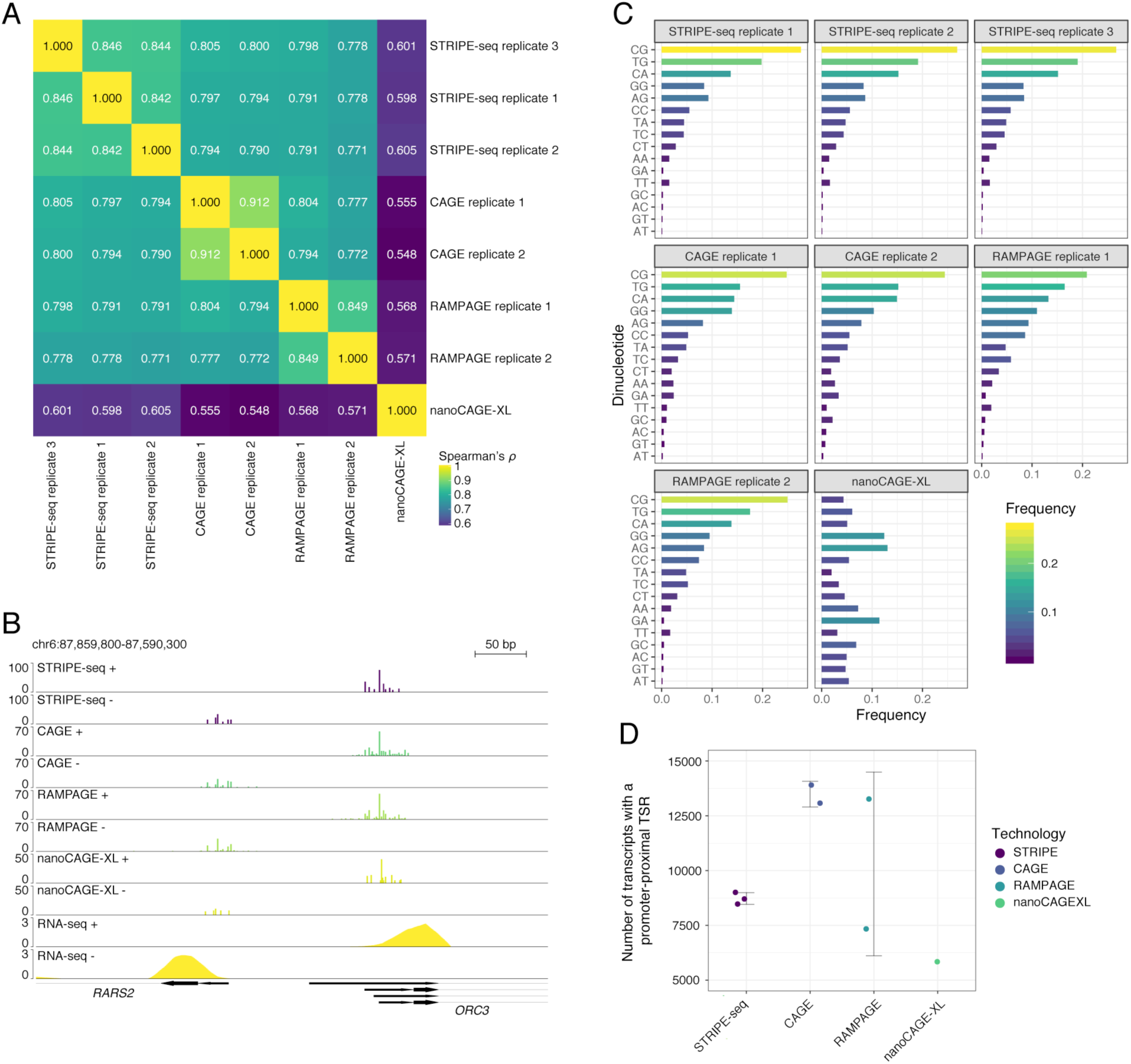
Comparison of human STRIPE-seq to CAGE, RAMPAGE, and nanoCAGE-XL. (A) Hierarchically clustered heatmap of Spearman’s *⍴* values for pairwise comparisons of STRIPE-seq, CAGE, RAMPAGE, and nanoCAGE-XL signal within promoter regions (−500 to +500 bp relative to an annotated TSSs). Prior to clustering, samples were thresholded such that each promoter had to have at least 3 read counts in one sample, and then counts were TMM normalized. (B) Genome browser-style tracks showing CPM-normalized STRIPE-seq, CAGE, RAMPAGE, nanoCAGE-XL (replicate 1 for each input amount), and poly(A)+ RNA-seq (replicate 1) at two representative regions of the human genome. (C) Dinucleotide frequencies at TSSs in all replicates of all technologies. (D) Jitter plot of the number of transcripts with a promoter-proximal TSR in each sample. Error bars represent standard deviation.

While the majority of TSSs detected by STRIPE-seq are promoter-proximal, a small fraction were located within genes or intergenic regions (Fig. 6C, Supplemental Figure 18A). Given that enhancers may be transcribed to generate enhancer RNAs (eRNAs) (Lam et al., 2014), we asked whether these distal TSSs were representative of eRNA initiation. To investigate this possibility, we first detected TSRs in K562 STRIPE-seq data and divided them into promoter-proximal (< 1 kb from an annotated TSS) and distal (≥ 1 kb from an annotated TSS) sets. Considering only TSRs present in all three STRIPE-seq replicates, we detected 6,648 proximal and 513 distal TRSs. The majority of distal TSRs detected by STRIPE-seq showed signal in CAGE, RAMPAGE, and nanoCAGE-XL datasets, suggesting that they are unlikely to be artifacts (Supplemental Figure 22A). We next compared proximal and distal TSRs to a set of 43,119 K562 enhancers from EnhancerAtlas 2.0, a compendium of enhancers generated by integrative analysis of a dozen genome-wide methods (Gao and Qian, 2019). Only 310 of 6,648 (4.7%) of proximal TSRs overlapped enhancers, while 179 of 513 (34.9%) distal TSRs coincided with annotated enhancers, a significantly higher proportion by Pearson’s chi-squared test (*p* = 9.49×10^−150^). Consistent with this, heatmap visualization of histone modification cleavage under targets and tagmentation (CUT&Tag) (Kaya-Okur et al., 2019) enrichment around distal TSRs revealed that a fraction of these sites are highly enriched for H3K4me1 and H3K27ac but not H3K4me3 (Supplemental Figure 22B), a characteristic signature of active enhancers (Creyghton et al., 2010; Heintzman et al., 2009; Rada-Iglesias et al., 2011; Zentner et al., 2011). These large-scale analyses were confirmed by visual inspection at specific loci. For instance, at a TSR within an intron of *LDLRAD3*, we observed signal in all analyzed TSS datasets as well as strong H3K4me1 and H3K27ac CUT&Tag signal (Supplemental Figure 22C). This active enhancer signature was also observed at an example intergenic site containing a distal TSR (Supplemental Figure 22C). We conclude that, even with modest input (100 ng total RNA), STRIPE-seq can detect a moderate number of eRNAs. We further note that the number of putative eRNA TSRs identified here is likely a conservative estimate, as the number of distal TSRs in total is 2,948 when a TSR is only required to be present in a single sample. The presence of many single-replicate distal TSRs may be due in part to the generally low abundance of eRNAs (De Santa et al., 2010; Rahman et al., 2017). Alternatively, some eRNA TSRs might be removed due to thresholding if they are infrequently transcribed and/or unstable.

## Discussion

Here we present STRIPE-seq, a rapid and efficient method for genome-wide profiling of TSSs. Requiring only a TSRT reaction and PCR amplification following depletion of uncapped RNA, STRIPE-seq is a simple and cost-effective protocol that can be performed in any molecular biology laboratory in less than 5 hours for $11.73 USD/sample. STRIPE-seq thus provides a substantial savings in terms of time and cost versus other TSS profiling methods (Supplemental Figure 23, Supplemental Table 4). STRIPE-seq encompasses a number of modifications to TSRT-based cDNA 5’ end profiling methods that address concerns related to efficiency and bias (Supplementary Figure 1). First, to reduce the occurence of spurious TSSs within gene bodies due to premature template switching, addition of the TSO is not added until reverse transcription has proceeded for 5 min. Second, to minimize TSO chaining, we rigorously tested TSO sequences for intrinsically low concatemerization potential and modified the 5’ end of the final TSO with biotin. As we were reproducibly unable to generate STRIPE-seq libraries from oligo- only samples (i.e. no-RNA samples) (Supplemental Figure 3), these strategies are effective in minimizing TSO concatemerization. Third, we used SPRI bead cleanups rather than tagmentation to optimize the size distribution of the cDNA library, removing the post-tagmentation PCR step necessary in nanoCAGE when ≥ 50 ng of total RNA are available, and thus minimizing the PCR cycles required for library amplification. In addition to the STRIPE-seq methodology, we provide an end-to-end computational workflow that enables reproducible analysis of STRIPE-seq data. With appropriate modifications to parameters at the alignment and processing steps, GoSTRIPES and TSRchitect can be used to analyze TSSs and TSRs derived from any related mapping technology. Furthermore, we are currently developing an R package, TSRexploreR, that will allow robust exploration of TSSs and TSRs derived from our method and processing workflow.

The major limitation of STRIPE-seq, and indeed any TSRT-based method, is the efficiency of TSRT itself (Wulf et al., 2019). While we were able to lower the number of PCR cycles required for library amplification relative to related methods using tagmentation or semi-suppressive PCR (Poulain et al., 2017), the frequencies of PCR duplicates in STRIPE-seq libraries is indicative of an upper limit on library complexity with the current iteration of the protocol. In our analysis of STRIPE-seq using increasing amounts of yeast total RNA, we observed a positive relationship between the proportion of unique read pairs retained and the amount of RNA used, indicating that library complexity is partially limited by input amount. This is not a practical limitation for STRIPE-seq in budding yeast and would presumably not complicate STRIPE-seq in other organisms with similar numbers of expressed transcripts, as our data indicate comparable sensitivity to both SLIC-CAGE and nanoCAGE. However, STRIPE-seq of more complex transcriptomes indicates loss of a few thousand transcripts relative to CAGE and a deeply sequenced RAMPAGE sample. Thus, in its present form, STRIPE-seq has reduced sensitivity in large transcriptomes relative to current high-input gold-standard methods at low to moderate sequencing depths. However, given the simplicity and rapidity of the protocol, its low cost, and modest input requirements, we think that it will be broadly useful and a practical approach for exploring transcription initiation on a global scale. Future development of STRIPE-seq will focus on improving TSRT efficiency to increase the complexity of the initial cDNA pool in order to overcome this problem. A recent rigorous examination of TSRT parameters (Wulf et al., 2019) may provide some improvements in this area. For instance, adding excess dCTP to a TSRT reaction was shown to almost double TSRT efficiency.

An exciting potential application of STRIPE-seq is TSS profiling in precious clinical or developmental samples where only modest quantities of RNA are available. We found that STRIPE-seq can reliably generate libraries from as little as 50 ng of total RNA. Assuming a total RNA content of 1-50 pg per mammalian cell (Han and Lillard, 2000; Islam et al., 2011; Livesey, 2003), STRIPE-seq could, in its present form, be used with the amount of RNA extracted from fewer than 50,000 such cells, depending on yield. Furthermore, we envision that, in conjunction with methodologies for single cell isolation and lysis such as those used in Smart-seq2 (Picelli et al., 2013), STRIPE-seq could be adapted for single-cell profiling of TSSs. The demonstrated sensitivity of STRIPE-seq for low-abundance and/or unstable transcripts such as eRNAs could also potentially be enhanced by using nascent rather than total RNA as input for library preparation. Furthermore, the addition of barcodes during the TSRT reaction opens the possibility of pooling samples after this step, allowing the processing of multiple STRIPE-seq samples at the same time to further save on cost and time. Lastly, given the simplicity of the STRIPE-seq method, we anticipate that some or all of the protocol could be automated to enhance throughput.

## Methods

### Biological samples

Yeast strain S288C was grown in YPD medium at 30°C under constant agitation. Where indicated, cells were treated with 1.5 mM diamide under the same growth conditions for 1 h. Yeast total RNA was extracted with the MasterPure Yeast RNA Purification Kit (Lucigen MPY03100) as per the manufacturer’s protocol. Human K562 cells were grown in DMEM + 10% FBS and 1x penicillin/streptomycin at 37°C with 5% CO_2_. K562 total RNA was extracted with TRIzol (Invitrogen) as per the manufacturer’s protocol, treated with DNase I, and purified using RNAClean XP beads (Beckman Coulter) at a beads:sample ratio of 1.8:1 as per the manufacturer’s protocol.

### STRIPE-seq

A step-by-step STRIPE-seq library construction protocol is available at protocols.io (https://www.protocols.io/view/stripe-seq-library-construction-2ivgce6). See Supplemental Table 5 for oligo sequences.

#### TEX treatment of total RNA

For TEX digestion, 50, 100, or 250 ng total RNA (≥ 25 ng/µL) was combined with 0.2 µL Terminator Reaction Buffer A (10X), 0.1 µL TEX (1 U/µL) and H_2_O to 2.0 µL. The reaction was then incubated at 30°C for 1 h. EDTA inhibits TEX and should thus be excluded from the RNA storage buffer. Due to the small volumes involved, we recommend making a master mix sufficient for the desired number of samples. We also note a higher proportion of rRNA reads in the 100 ng and especially the 250 ng samples, and thus suggest using 0.2 µL of TEX above 100 ng of RNA input.

#### Template-switching reverse transcription

Template-switching reverse transcription (TSRT) was performed as previously described in Smart-seq2 (Picelli et al., 2013) with the following modifications. The initial primer mix was prepared by combining 1 µL RTO (10 µM) with a different barcode per sample (see Supplemental Table 5 for barcode sequences and https://github.com/rpolicastro/barcode-generator for a barcode generation script used here), 0.5 µL dNTP mix (10 mM each dNTP), and 1.5 µL of a mixture of sorbitol (6.6 M) and trehalose (0.3 M) prepared as previously described (Batut and Gingeras, 2013). The 2 µL TEX digestion was added to the primer mix, and the solution was incubated at 65°C for 5 min, 4°C for 2 min and then immediately placed on ice. 5 µL of a reverse transcription master mix containing 2 µL SuperScript II First Strand Buffer (5X), 0.5 µL DTT (0.1 M), 2 µL betaine (5 M), and 0.5 µL (100 U) SuperScript II reverse transcriptase (200 U/µL) was added to the RNA mixture. For reverse transcription (RT), the sample was incubated at 25°C for 10 min and 42°C for 5 min. 0.25 µL of TSO (400 µM) was added to the reactions while still in the heat block and then mixed thoroughly by pipetting. The reaction was completed with a 25 min incubation at 42°C and a 10 min incubation at 70°C. cDNA was purified using 8 µL of RNAclean XP beads (0.8:1 beads:sample ratio) according to the manufacturer’s protocol. For both this and subsequent cleanup steps, AMPure XP (Beckman Coulter) or analogous SPRI bead preparations may be substituted to save on costs. First-strand cDNA was eluted in 12 µL nuclease-free H_2_O.

#### Second-strand cDNA synthesis/library PCR

Second-strand cDNA synthesis and library amplification was performed by combining 11 µL eluted first-strand cDNA with 12.5 µL of KAPA HiFi HotStart ReadyMix (2X; Kapa Biosystems, Inc.) and 0.75 µL each of the forward (10 µM) and reverse (10 µM) STRIPE-seq library oligos for a total reaction volume of 25 µL. The PCR mixture was thermal cycled using the following program: 95°C for 3 min, 16-20 cycles of 98°C for 20 s, 63°C for 15 s, and 72°C for 45 s, and a final extension at 72°C for 2 min. A two-sided size selection was then performed. First, to remove small fragments, 16.3 µL RNAClean XP beads were added (0.652 beads:sample ratio) and cDNA was purified according to the manufacturer’s protocol. cDNA was eluted in 17 µL nuclease-free H_2_O. To remove large fragments, 15 µL of the previous eluted cDNA were transferred to a new tube, 8.3 µL RNAClean XP beads were added (0.553:1 beads:sample ratio), and the bead/sample mixture was incubated at room temperature for 10 min. Beads were collected on a magnetic rack for 10 min and 22 µL supernatant was transferred to a new tube. To isolate the final cleaned cDNA library, 22 µL RNAClean XP beads (1:1 beads:sample ratio) were added and purification was completed as per the manufacturer’s protocol. The final library was eluted in 16 µL nuclease-free H_2_O and 15 µL was transferred to a new tube. Typical yields from the STRIPE-seq library protocol ranged from 25 to 100 ng of total library with a size distribution from 200-1000 bp as analyzed on an Agilent 2200 TapeStation instrument using a High Sensitivity D5000 ScreenTape device (Agilent Technologies).

#### Sequencing of STRIPE-seq Libraries

Libraries were pooled by molarity based on quantification of fragments between 180-900 bp as determined by the aforementioned TapeStation analysis. An additional round of RNAClean XP size selection (1:1 beads:sample ratio) was performed on the pooled library as needed to remove any residual putative primer-dimers ≤150 bp. Pooled libraries were denatured following the standard NextSeq procedure using a 1 nM initial library concentration. Libraries were sequenced for 75 cycles for both the R1 and R2 reads in paired-end mode on the Illumina NextSeq 500 platform at the Indiana University Center for Genomics and Bioinformatics (CGB). Note that, due to the homogeneity present in the TSO following the UMI (TATAGGG), a modest proportion of a STRIPE-seq run (20-30%) should be comprised of more random sequences either in the form of a spike-in such as PhiX or another type of library such as RNA-seq or ChIP-seq. We note that this caveat only applies to newer Illumina platforms (e.g. NextSeq and NovaSeq) using two-color chemistry, wherein G nucleotides are represented as the absence of color, thus leading to poor cluster acquisition and potential run failure when the TATAGGG sequence in the adapter is reached. This problem is less pronounced on four-color Illumina platforms (e.g. HiSeq and MiSeq) and so in these cases the proportion of spike-in and/or other library types can be decreased to 10-15%.

### RNA-seq

Total RNA (10 µg) was used to generate poly(A)+ libraries with the Illumina TruSeq Stranded mRNA Library Prep kit. Libraries were prepared by the CGB and sequenced as for STRIPE-seq samples.

### Data analysis

#### STRIPE-seq

##### Read processing and alignment

STRIPE-seq read files were processed and aligned to the respective genomes using the GoSTRIPES workflow (https://github.com/BrendelGroup/GoSTRIPES). The workflow is implemented as a GNU Make (https://www.gnu.org/software/make/) file, consisting of a section that defines program parameters, rules that define intermediate and final target outputs, and recipes for creating the targets. GoSTRIPES performs the following steps (shown as a dependency graph in Supplemental Figure 4):

1. (Optional) Analysis of read quality (FastQC (Andrews, 2010))
2. Trimming of adapter sequences (Trimmomatic (Bolger et al., 2014))
3. Removal of reads mapping to rRNA species (TagDust2 (Lassmann, 2015))
4. Selection of reads with the UMI-TATAGGG sequence (*selectReadsByPattern.pl*)
5. Addition of UMI sequence to read name, to facilitate PCR duplicate removal in single-end sequencing if necessary (UMI-tools (Smith et al., 2017))
6. Trimming of adapter and TATAGGG sequences (Cutadapt (Martin, 2011))
7. Rescue of a few remaining reads with reliable detection of template switching chaining (*selectReadsByPattern.pl*)
8. (Optional) Analysis of processed read quality post cleanup (FastQC (Andrews, 2010))
9. Alignment to reference genome (STAR (Dobin et al., 2013))
10. Removal of duplicate reads based on matching R1 and R2 read positions for paired-end samples (SAMtools) (Li et al., 2009). For single-end sequencing, the UMI previously stored in the read name maxy be used to remove PCR duplicates.
11. Scrubbing of aligned reads to remove reads with excessive soft clipping, very short or very long within-read gaps, and pairs with excessively long template lengths (*scrubSAMfile.pl*)

All programs and dependencies required to complete this workflow are available in the GoSTRIPES Singularity container (Kurtzer et al., 2017) which can be downloaded from Singularity Hub (Sochat et al., 2017) (https://www.singularity-hub.org/collections/1204). Parameter settings used are accessible as described in the next section.

##### Identification of TSSs and TSRs

Starting from the alignments of GoSTRIPES-processed STRIPE-seq libraries in BAM format, we used the R Bioconductor package TSRchitect (https://bioconductor.org/packages/release/bioc/html/TSRchitect.html) (Raborn et al., 2017) to identify TSSs and cluster them into TSRs. At the conclusion of the analysis with TSRchitect we exported TSS and TSR datasets to text-delimited files, including BED format. The output of TSRchitect can also be used with the new Bioconductor package CAGEfightR (Thodberg et al., 2019) for analyses not described here, such as discovery and association of putative enhancers with target genes. As with GoSTRIPES, the TSRchitect package with all its dependencies can be run via Singularity, which greatly facilitates reproducibility on any UNIX-based system. The entirety of our workflows consisting of GoSTRIPES-enabled read processing and alignment and TSRchitect-enabled identification of TSSs and TSRs are documented for both the yeast and human data at our Github site TSRexplore (https://github.com/BrendelGroup/TSRexplore). With just a few minor edits to configuration scripts, a reader will be able to reproduce all of the data analysis work from this paper. Moreover, the provided scripts can easily be modified to allow analysis of other TSS data sets, independent of the methodology used.

##### Downstream analysis

To facilitate straightforward and reproducible downstream analysis of STRIPE-seq data, we developed a series of integrated scripts that will form the basis of an R package, TSRexploreR (manuscript in preparation). In this paper, TSRexploreR was used to explore TSS read thresholds, determine the genomic distributions of TSSs, assess TSS and TSR correlation between samples, determine TSS density relative to annotated start codons and promoters, analyze the sequence context of TSSs, perform differential TSR and expression analysis, and perform GO analysis. The version of TSRexploreR used here is available at https://github.com/zentnerlab/tsrexplorer/tree/policastro_etal_2020. For analysis of yeast samples, we used the Ensembl 98 R64-1-1 yeast genome sequence (Saccharomyces_cerevisiae.R64-1-1.dna.toplevel.fa) and annotation file (Saccharomyces_cerevisiae.R64-1-1.98.gtf). For analysis of K562 samples, we used the Ensembl 98 GRCh38.p13 soft-masked human genome sequence (Homo_sapiens.GRCh38.dna_sm.primary_assembly.fa) and annotation file (Homo_sapiens.GRCh38.98.chr.gtf). Scripts used to automate TSRexploreR functions used in this work (*yeast_TSRexploreR.R*, *human_TSR_exploreR.R*) are available at https://github.com/zentnerlab/stripeseq/tree/policastro_etal_2020. For visualization of TSS mapping data, we used Gviz (Hahne and Ivanek, 2016) with custom automation scripts (*yeast_Gviz.R*, *human_Gviz.R*), available at https://github.com/zentnerlab/stripeseq/tree/policastro_etal_2020.

##### Promoter correlation analysis

To generate sets of protein-coding gene promoters, we read the GTFs listed above into R as TxDb objects and subjected them to a filtering step. For yeast, we required that the transcript name contain “mRNA” and excluded mitochondrial transcripts, yielding a set of 6,572 transcripts. For human, we required that the transcript name be present in a list of 168,608 protein-coding transcripts obtained from Ensembl and excluded transcripts originating from the Y or mitochondrial chromosomes, yielding a set of 152,701 transcripts. Promoter windows (−250 to +100 for yeast, −500 to +500 for human) were obtained from the corresponding TxDb object and filtered against the lists of mRNA IDs generated above. We then counted the number of TSSs from each sample within each promoter window, normalized counts, and performed Spearman correlation analysis. Scripts used for promoter correlation analysis (*yeast_promoter_correlation.R*, *human_promoter_correlation.R*) are available at https://github.com/zentnerlab/stripeseq/tree/policastro_etal_2020.

#### RNA-seq

rRNA reads were first removed from paired-end FASTQ files using TagDust2 (Lassmann, 2015) (v2.33.0) with additional settings -fe 3 -dust 97. STAR (Dobin et al., 2013) (v2.7.0e) was then used to generate genome indices (with the additional setting *--genomeSAindexNbases 10* for yeast) and align reads to the GRCh38.p13 or R64-1-1 genome assemblies for human and yeast data, respectively. Finally, the aligned reads were counted to the nearest overlapping feature using the Subread (Liao et al., 2019) (v1.6.4) function *featureCounts* with the settings *-t exon –g gene_id --minOverlap 10 --largestOverlap -s 2 -p -B*. bigWig files representing RNA-seq coverage were generated with deepTools (Ramírez et al., 2016) (v3.3.0) as follows: bamCoverage *--normalizeUsing CPM -bs 1 --filterRNAstrand [forward or reverse] -- smoothLength 25*. RNA-seq libraries were constructed using the deoxy-UTP (dUTP) method, so reads corresponding to positively-stranded transcripts were obtained with *--filterRNAstrand forward*, which excludes positively-stranded reads, as negatively-stranded reads correspond to positively-stranded transcripts in the dUTP protocol. Similarly, reads corresponding to negatively-stranded transcripts were obtained with *--filterRNAstrand reverse*. Reads were not extended so as not to generate coverage of skipped regions (introns). STRIPE-seq fragments in transcripts were counted with Subread featureCounts as above. Differential expression analysis was performed with edgeR (Robinson et al., 2010) as part of TSRexploreR.

#### Distal TSR analysis

##### Overlap of TSRs with enhancers

TSRs from three K562 100 ng STRIPE-seq replicates were merged to yield a set of 16,745 TSRs and annotated with ChIPseeker (Yu et al., 2015). Only TSRs with signal in all three replicates (n = 7,161) were considered for the ensuing analysis. We considered TSRs < 1 kb from an annotated TSS to be proximal and TSRs ≥ 1 kb from a TSS to be distal. A set of 43,148 K562 enhancer annotations (using hg19 coordinates) were obtained from EnhancerAtlas 2.0 (Gao and Qian, 2019) (http://www.enhanceratlas.org/data/download/enhancer/hs/K562.bed) and converted to hg38 coordinates with UCSC liftOver, yielding a set of 43,119 enhancers. Proximal and distal TSRs were then overlapped with enhancer coordinates and counted. Proportions of proximal and distal TSRs overlapping enhancers were compared using Pearson’s chi-squared test. The script used for this analysis (*distal_TSR_analysis.R*) is available at https://github.com/zentnerlab/stripeseq/tree/policastro_etal_2020. To enable display of enhancer annotations in Gviz, chromosome names were converted from UCSC to Ensembl format (i.e. chr1 to 1) with sed ‘/s/chr//g’, as we used Ensembl genome sequences and annotations for STRIPE-seq analysis.

##### Analysis of TSS signal at distal TSRs

CPM-normalized TSS bedGraphs were converted to bigWigs with UCSC bedGraphToBigWig to enable compatibility with deepTools (Ramírez et al., 2016) v3.3.0. Signal around TSR midpoints was then determined with deepTools *computeMatrix* and heatmaps were generated with the deepTools function *plotHeatmap*, sorting all heatmaps descending by mean STRIPE-seq minus strand signal. The nearly exact splitting of the heatmap into plus and minus strand signal likely arises from the fact that most TSRs only display signal on one strand in each method at these TSRs.

##### Analysis of histone modifications at distal TSRs

K562 CUT&Tag datasets (Kaya-Okur et al., 2019) were downloaded from the SRA, converted to FASTQ format with fastq-dump (Leinonen et al., 2010), and aligned to the GRCh38 and Ensembl EB1 (*E. coli*) genome assemblies with Bowtie2 (Langmead and Salzberg, 2012) v2.3.5 with default parameters plus *-I 10 -X 700 --no-unal --no-discordant --no-mixed --dovetail*. Replicate BAM files were merged with SAMtools and a normalization factor corresponding to (1,000/number of fragments mapped to EB1) was calculated for each sample. Spike-in-normalized bigWig signal tracks were generated with deepTools *bamCoverage* with the *-- scaleFactor* flag set to the appropriate normalization factor and *-bs 25 -e* to set the resolution to 25 bp and extend reads to their paired-end fragment length. Signal around TSR midpoints was then determined with deepTools *computeMatrix* and plotted as heatmaps with deepTools *plotHeatmap*, sorting all heatmaps in descending order by mean H3K4me1 signal. For the purpose of displaying CUT&Tag bigWig files with Gviz, chromosome names were converted from UCSC to Ensembl format using *convertBigWigChroms.py* (https://gist.github.com/dpryan79) with the *GRCh38_UCSC2ensembl* conversion file (https://github.com/dpryan79/ChromosomeMappings). Scripts used for generation of TSS and CUT&Tag bigWigs (*bedgraphtobigwig.sh*, *merge_ecoli.sh*, *merge_human.sh*) and heatmaps (*distal_TSR_heatmaps.sh*) are available at https://github.com/zentnerlab/stripeseq/tree/policastro_etal_2020.

#### Public datasets

All raw datasets were obtained from the SRA using the accession numbers listed in Supplemental Table 6. Yeast Ribo-seq WIG files were obtained from GEO (GSM1949550/1) (Nissley et al., 2016). Chromosome names in the Ribo-seq WIG files were converted from UCSC to Ensembl format (e.g. chrI to I) with sed *s/=chr/=/g* for display in Gviz.

#### Data availability

Raw STRIPE-seq and RNA-seq data are available from the SRA (SRP238585) and processed data are available from GEO (GSE142524).

## Acknowledgements

We thank Jie Huang, Nathan Keith, and David Miller for assistance with development and testing of the STRIPE-seq library construction protocol and Sungyun Kang for K562 cell culture. This work was supported by the Indiana Clinical and Translational Sciences Institute, funded in part by grant UL1 TR001108 from the National Institutes of Health, National Center for Advancing Translational Sciences, Clinical and Translational Sciences Award to R.T.R., National Science Foundation grant IOS-1221984 to V.P.B., and Indiana University startup funds and National Institutes of Health grant R35GM128631 to G.E.Z.

## Author contributions

R.A.P. developed the method and performed all experiments. R.A.P., V.P.B., and G.E.Z. analyzed the data. R.A.P., R.T.R., and V.P.B. wrote software. R.A.P., R.T.R., V.P.B., and G.E.Z. interpreted the results and wrote the manuscript.

