## Supplemental material (Supplemental Tables 1-6 legends, Supplemental Figures 1-23) for "Simple and efficient measurement of transcription initiation and transcript levels with STRIPE-seq"

Contents:

Legends for Supplemental Tables 1-6

Supplemental Figures 1-23

Supplemental References

**Supplemental Table 1**

Processing and alignment information for yeast and human STRIPE-seq samples

**Supplemental Table 2**

Information on TSRs differentially regulated by diamide treatment of yeast.

**Supplemental Table 3**

Differential expression data from STRIPE-seq and RNA-seq measurement of transcript abundances in control and diamide-treated yeast.

**Supplemental Table 4**

Cost and time analysis of TSS mapping methods. Primers, reagent lists, and protocols for SLIC-CAGE (Cvetesic et al., 2018), nanoCAGE (Poulain et al., 2017), nAnT-iCAGE (Murata et al., 2014), and RAMPAGE (Batut and Gingeras, 2013) were retrieved from published protocols. Reagent costs were calculated based on price without institutional discounts. Costs exclude common laboratory chemicals such as NaCl and NaOH, and reagents with a negligible per sample cost were not considered in per sample calculations. Number of steps and estimated timings were based on the above protocols, taking into consideration the published time estimates.

**Supplemental Table 5**

Oligonucleotide sequences used for STRIPE-seq library construction.

**Supplemental Table 6**

Previously published datasets used in this work.

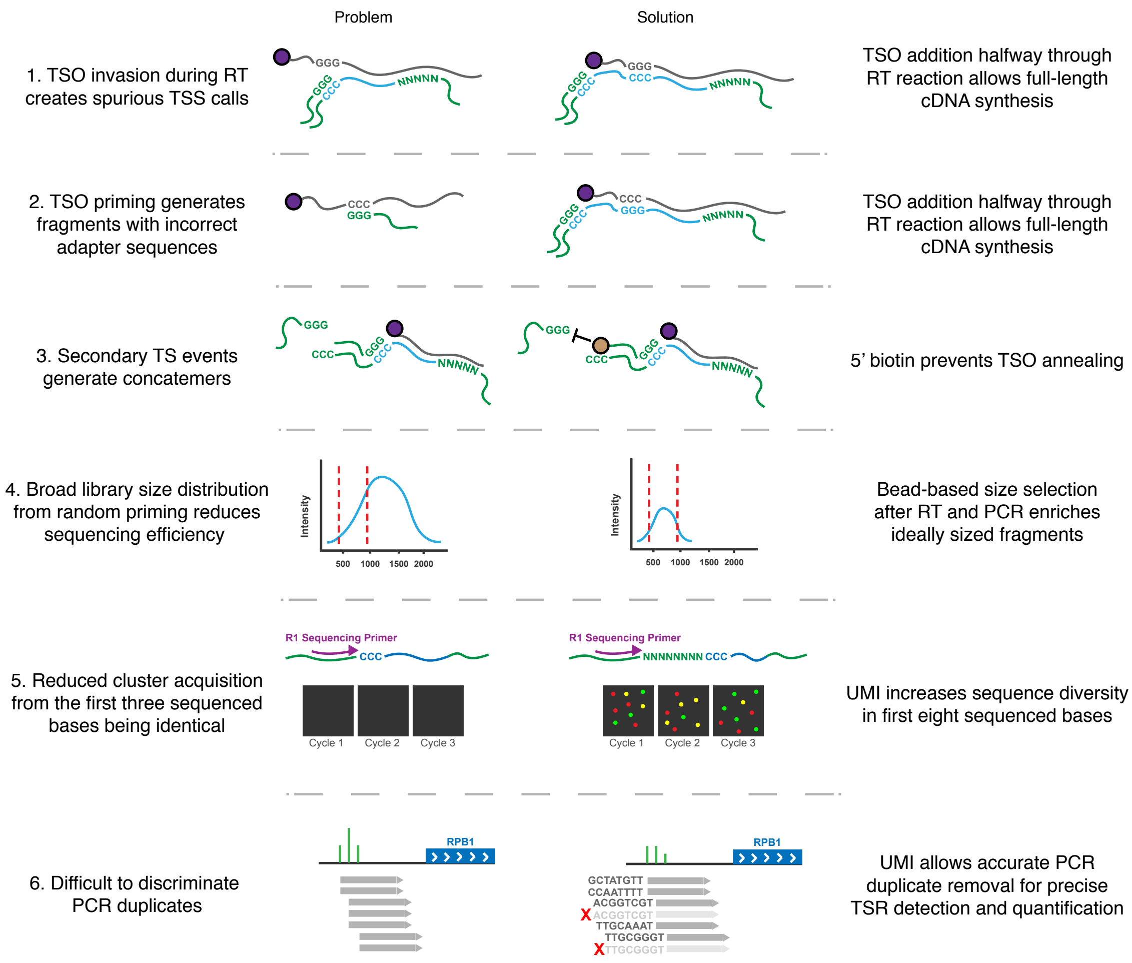


**Supplemental Figure 1. Steps taken to mitigate drawbacks of current TSRT-based TSS profiling methods**

1. During TSRT, the TSO can hybridize to poly(C) stretches in the nascent cDNA transcript (TSO invasion), leading to artifactual TSSs within gene bodies. Addition of the TSO 5 minutes into the reaction ensures that a majority of cDNA transcripts are fully extended to help suppress this artifact. 2) The TSO can also act as a primer, annealing to CCC sequences in RNA molecules. This is also suppressed by later addition of the TSO to the TSRT reaction.3) Additional TSOs can hybridize to a CCC tract added to the end of the first TSO by reverse transcriptase. This phenomenon is reduced by modification of the 5' end of the TSO with a biotin moiety. 4) TSRT produces a wide range of cDNA sizes, with very large and very small molecules incompatible with Illumina sequencing. Tagmentation with Tn5 transposase is frequently used to optimize library size, but requires an additional PCR step. In STRIPE-seq, SPRI bead-based size selection is used to optimize library size. 5) On Illumina sequencers that use two-color chemistry (e.g. NextSeq, NovaSeq), G is represented as the absence of color. The

homogeneity inherent to the TSO (TATAGGG) can lead to reduced cluster acquisition and loss of data. To improve sequence diversity, the STRIPE-seq TSO has an 8-nt unique molecular identified (UMI) as the first 8 bases sequenced. 6) The presence of the UMI also allows for removal of PCR duplicates when single-end sequencing is used, providing more accurate quantification of TSSs.

Reverse transcription oligo (RTO)

5’ CAAGCAGAAGACGGCATACGAGAT[i6 Barcode]GTGACTGGAGTTCAGACGTGTGCTCTTCCGATCT NNNNN 3’

| | |
|---|---|
| Full Illumina P7 adapter | Theoretical sample pooling after RT |
| Barcode | TruSeq multiplexing for sequencing |
| Random pentamer | Reverse transcription and reduced transcript length bias |

Template-switching oligo (TSO)

5’ Biotin CCTACACGACGCTCTTCCGATCT NNNNNNNN TATA rGrGrG 3’

| | |
|---|---|
| Biotin | Reduced TSO chaining |
| Partial Illumina P5 adapter | Increased RT efficiency |
| Unique Molecular Identifier | Increased clustering efficiency and duplicate removal |
| Spacer | Reduced TSO invasion |
| Riboguanosines | Template switching |

**Supplemental Figure 2. Design of the STRIPE-seq TSO and RTO**

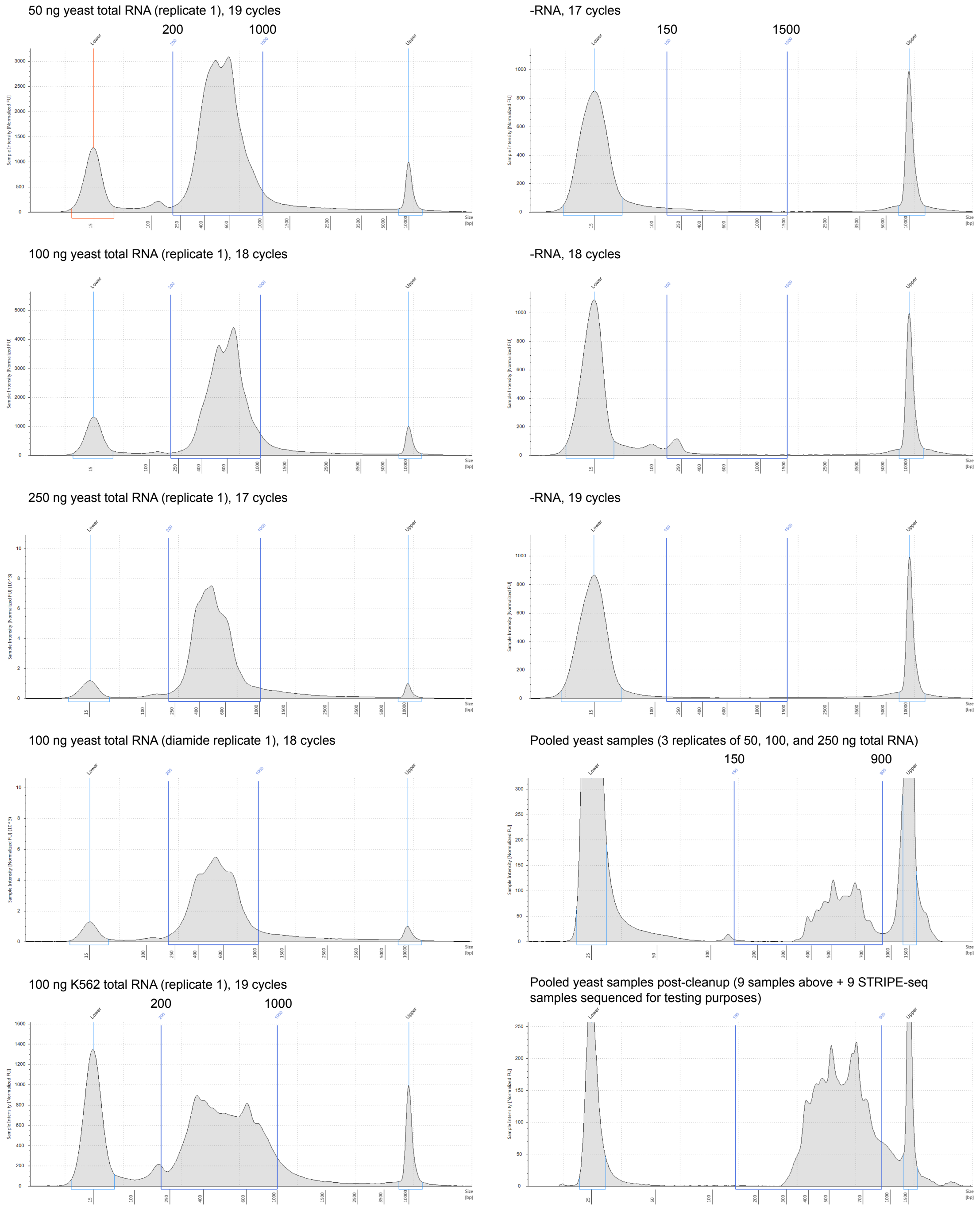


**Supplemental Figure 3. TapeStation analysis of STRIPE-seq libraries**

Shown are representative results for libraries constructed with 50, 100, and 250 ng of control yeast total RNA, 100 ng diamide-treated yeast total RNA, 100 ng K562 total RNA, no RNA input with the indicated number of PCR cycles, and pooled libraries before and after a mild bead cleanup (1:1 beads:sample ratio) to remove oligo dimers. Note that the trace shown for the pre-cleanup pool is for the 9 untreated yeast samples, while the trace shown for the post-cleanup pool is for these 9 samples plus 9 additional yeast samples sequenced for testing purposes but not presented here. Unpooled libraries were run on HSD5000 ScreenTapes and pooled libraries were run on HSD1000 ScreenTapes.

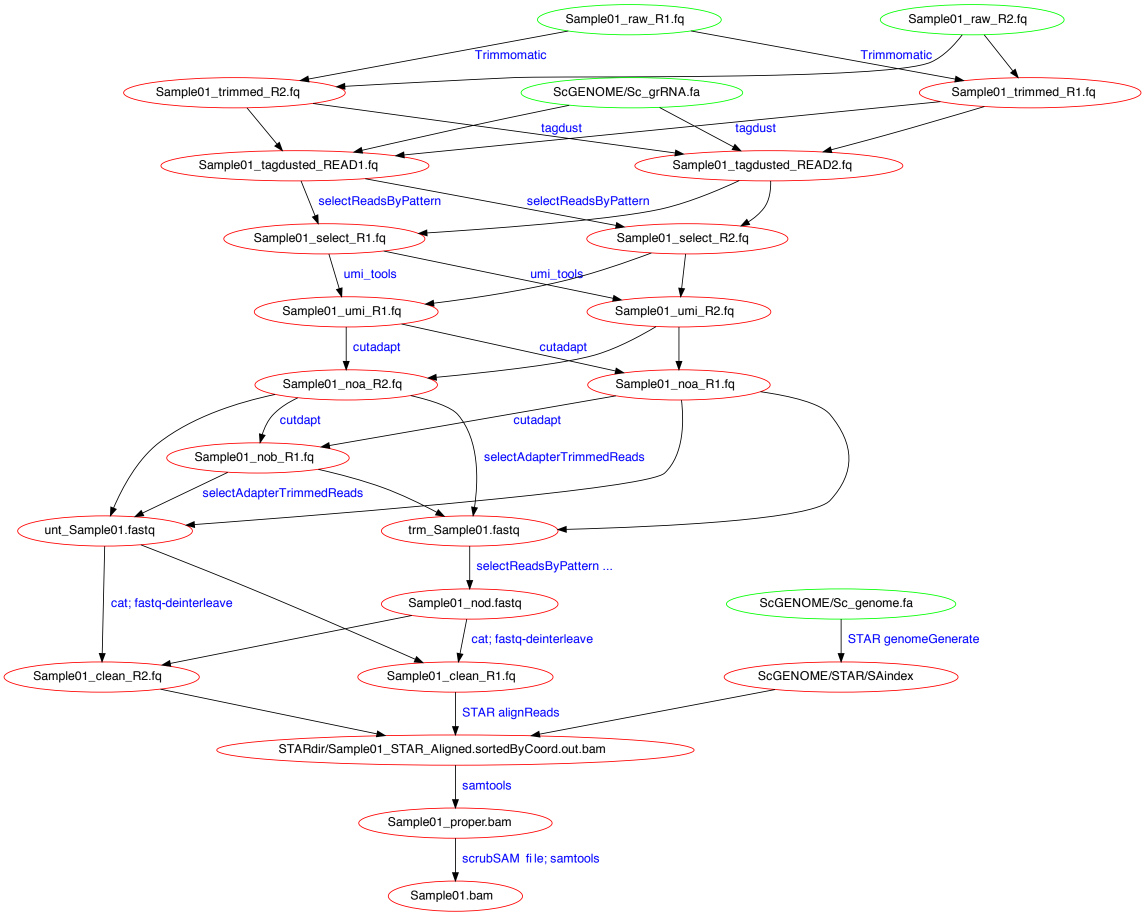


**Supplemental Figure 4. Dependency map of the GoSTRIPES computational workflow**

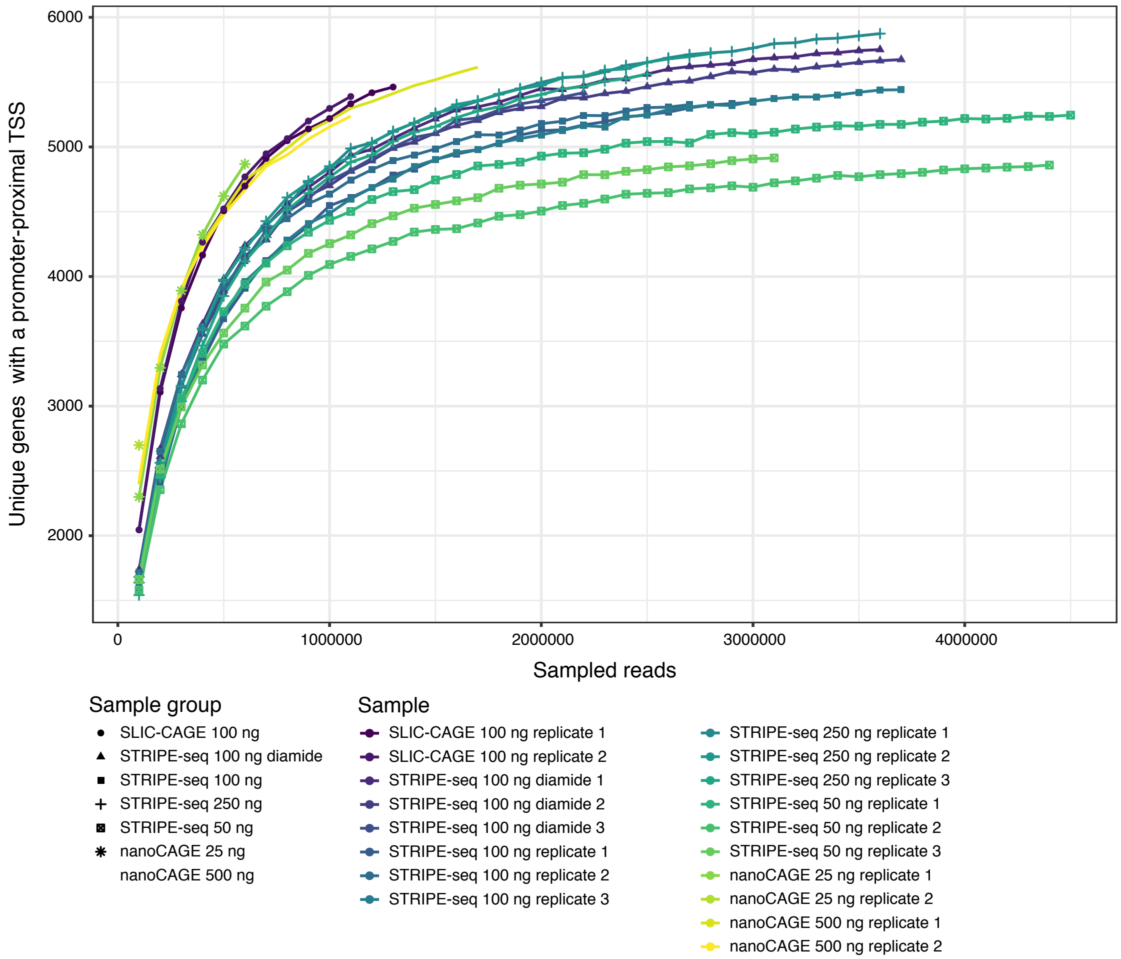


**Supplemental Figure 5. Saturation analysis of yeast TSS mapping datasets**

Analysis of sequencing saturation in STRIPE-seq, SLIC-CAGE, nanoCAGE. rRNA was computationally removed from FASTQ files, and then reads were aligned to the yeast R64-1-1 genome assembly. The given numbers of mappable reads were sampled from the BAMs, PCR duplicates were removed from STRIPE-seq samples, and only primary alignments with properly paired mates were retained. TSSs were then called using TSRchitect and annotated using ChIPseeker. Note that PCR duplicates could not be removed from SLIC-CAGE and nanoCAGE samples because the FASTQ files were single-end and, in the case of nanoCAGE, deposited with the UMI removed.

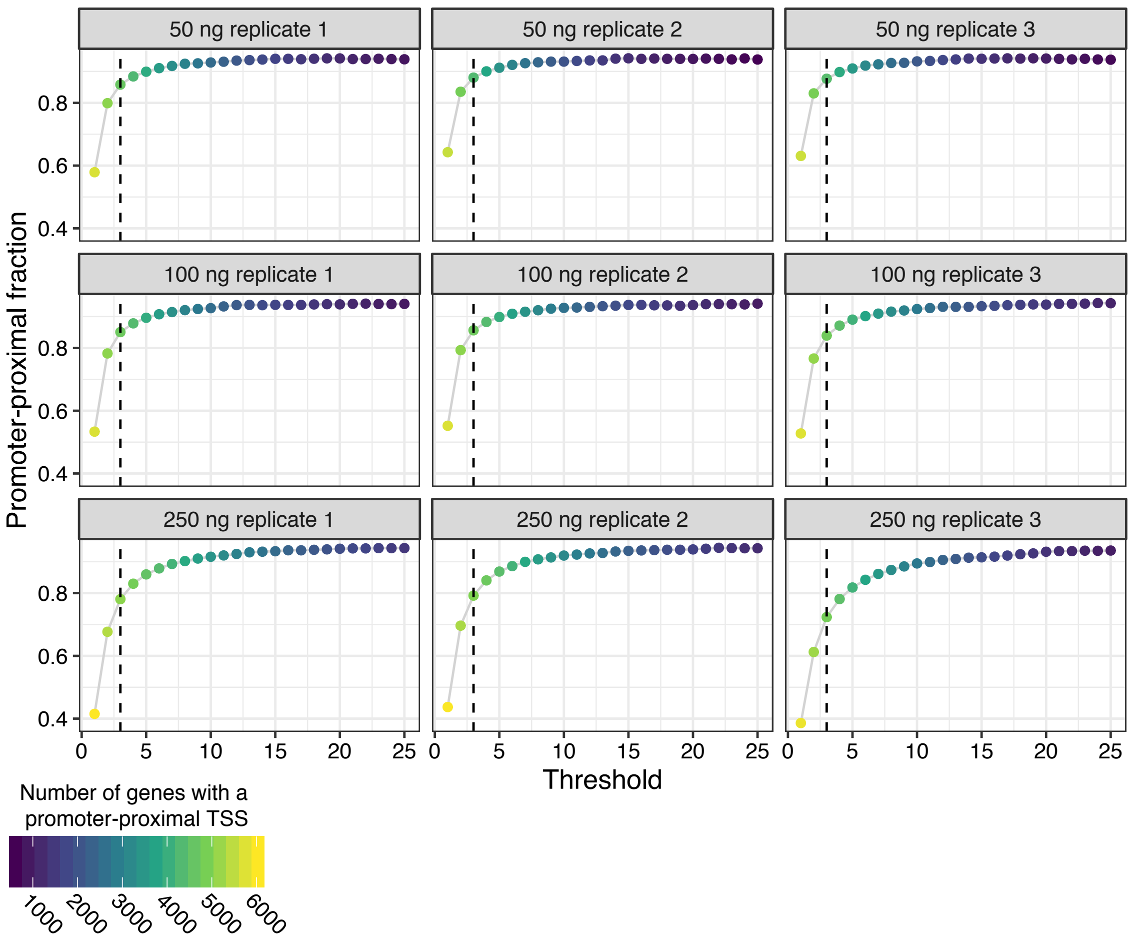


**Supplemental Figure 6. Threshold analysis of all yeast STRIPE-seq samples**

As in Figure 2A, plots show the fraction of unique TSSs that is promoter-proximal (-250 to +100 bp relative to an annotated start codon) at the indicated read threshold for each sample. Dot color indicates the number of genes with a promoter-proximal TSS.

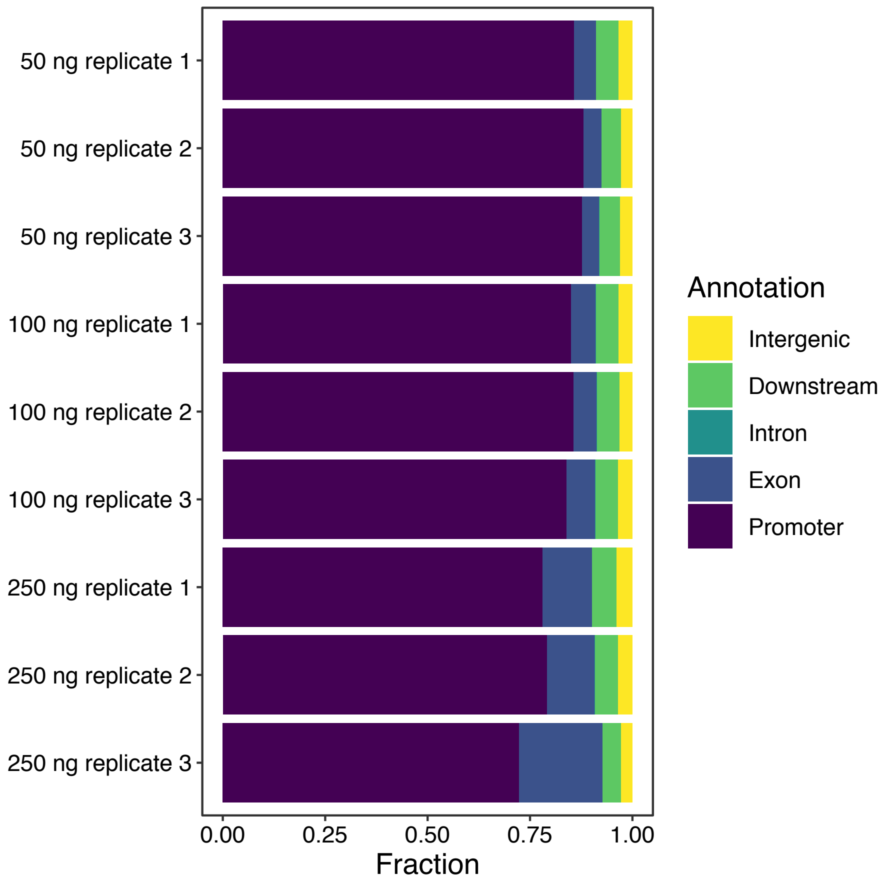


**Supplemental Figure 7. Genomic distribution of yeast TSSs identified by STRIPE-seq**

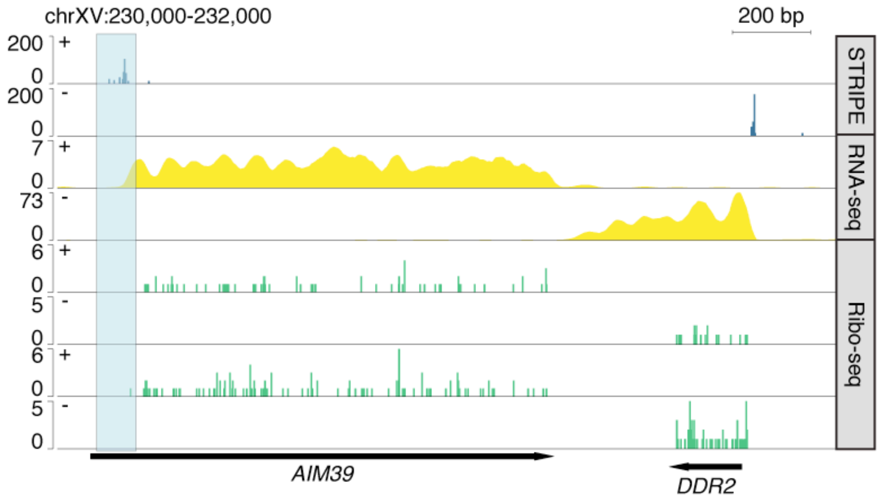


**Supplemental Figure 8. Analysis of ribosome profiling at *AIM39***

Genome browser-style tracks showing CPM-normalized STRIPE-seq and poly(A)+ RNA-seq as well as two replicates of Ribo-seq signal at the *AIM39* locus. STRIPE-seq signal (highlighted in blue) is upstream of the start of the RNA-seq signal, which in turn is upstream of the start of the Ribo-seq signal, strongly suggesting misannotation of the *AIM39* start codon.

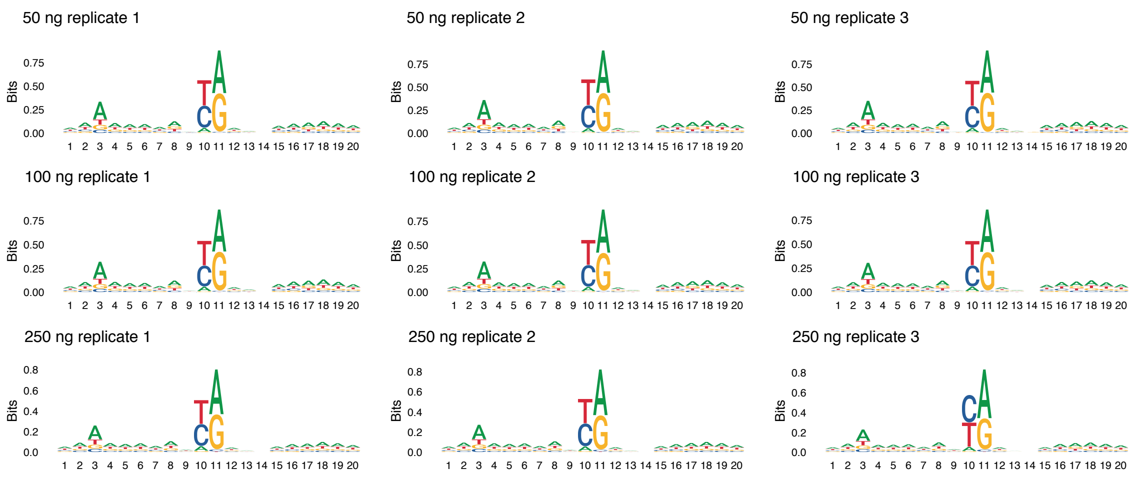


**Supplemental Figure 9. Sequence logos for yeast TSSs identified by STRIPE-seq**

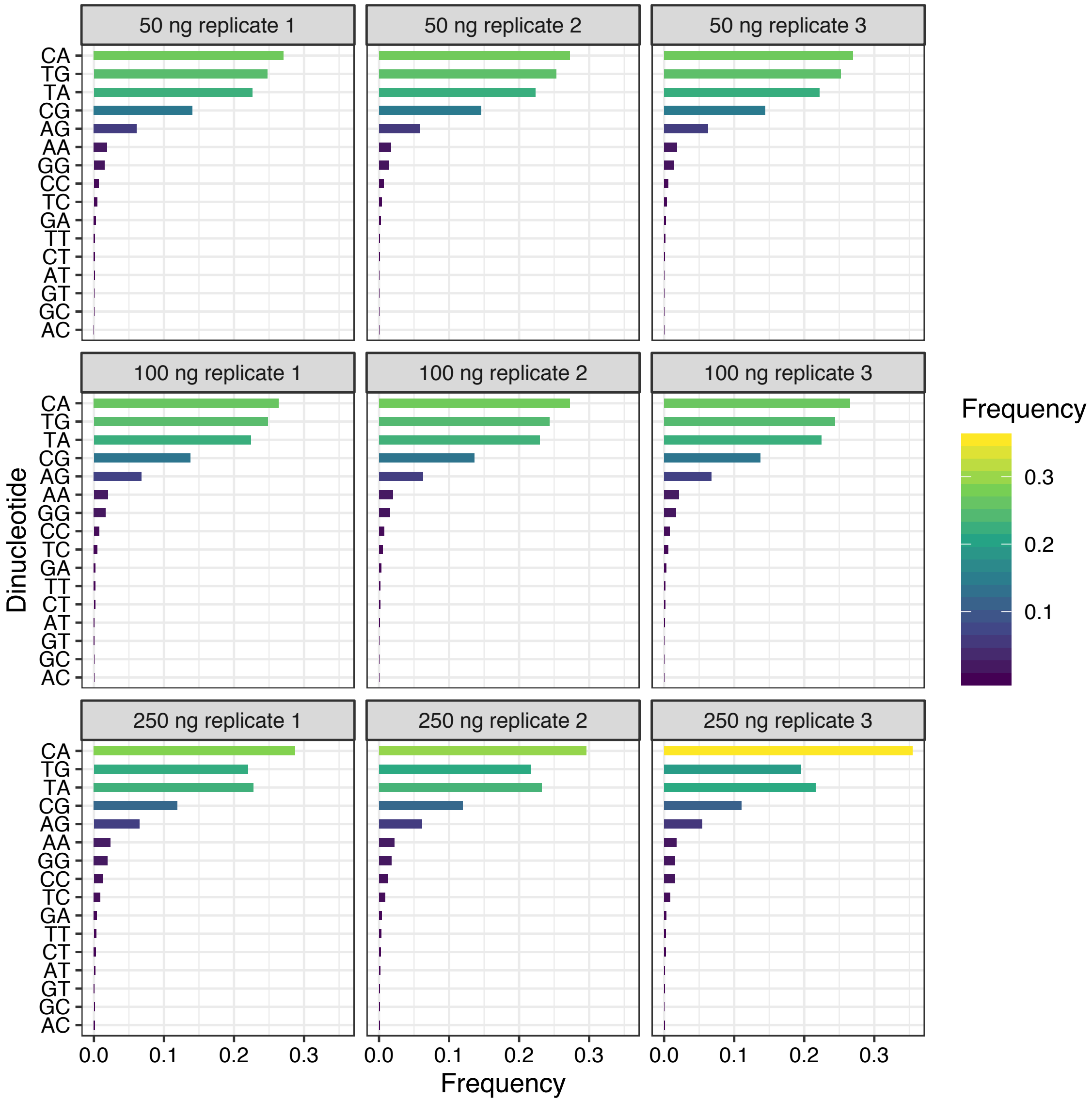


**Supplemental Figure 10. Dinucleotide frequencies at yeast TSSs identified by STRIPE-seq**

Dinucleotides are in descending order based on their average frequency across all samples.

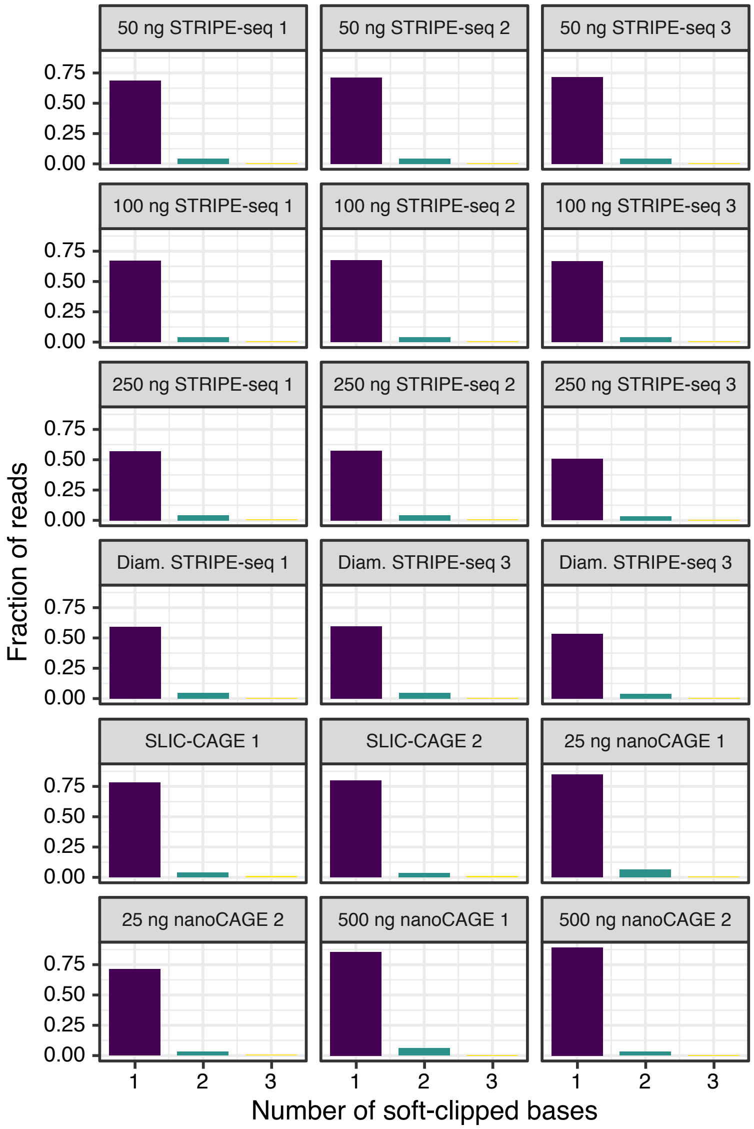


**Supplemental Figure 11. Analysis of soft-clipped base prevalence in yeast TSS mapping datasets**

Bar charts showing the frequencies of STRIPE-seq, SLIC-CAGE, and nanoCAGE R1 reads with the given number of soft-clipped bases extending upstream from the called TSS. Reads with more than 3 soft clipped-bases were infrequent in all methods and were thus not analyzed further.

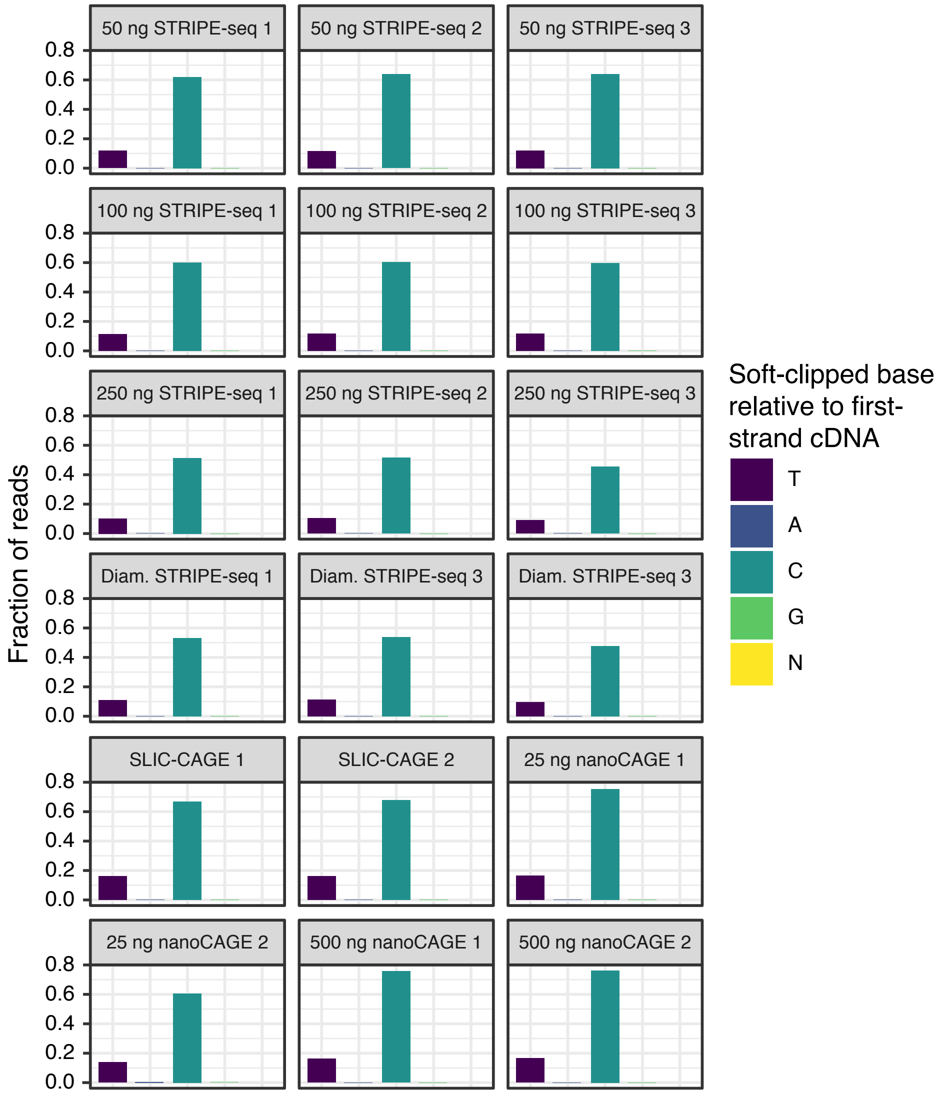


**Supplemental Figure 12. Sequence distribution of soft-clipped bases in yeast TSS mapping datasets**

Bar charts showing the frequencies of soft-clipped bases in STRIPE-seq, SLIC-CAGE, and nanoCAGE at the position directly upstream of the called TSSs. Bases displayed are relative to the first-strand cDNA.

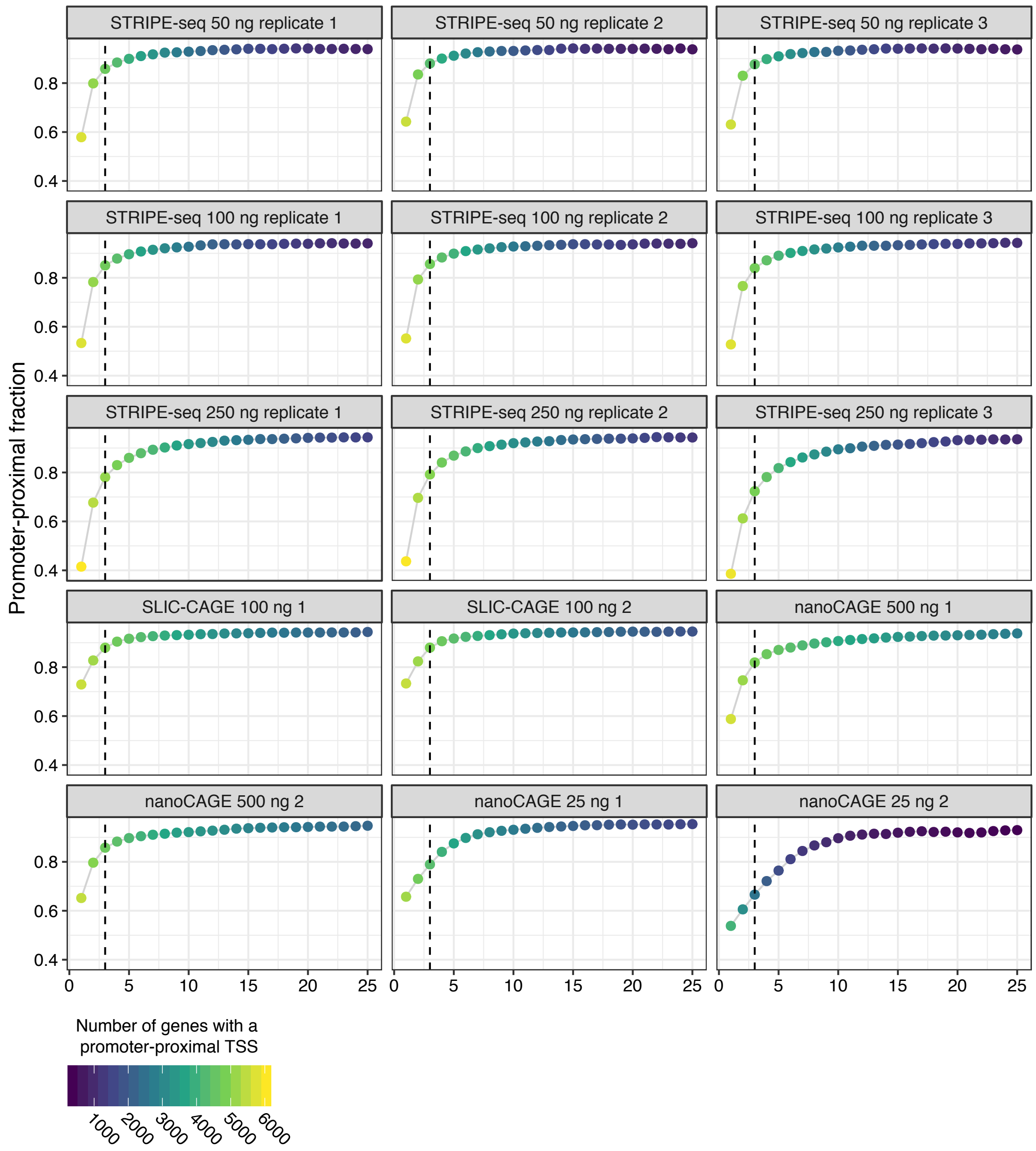


**Supplemental Figure 13. Threshold analysis of yeast STRIPE-seq, SLIC-CAGE, and nanoCAGE datasets**

As in Figure 2A, plots show the fraction of unique TSSs that is promoter-proximal (-250 to +100 bp relative to an annotated start codon) at the indicated read threshold for each sample. Dot color indicates the number of genes with a promoter-proximal TSS.

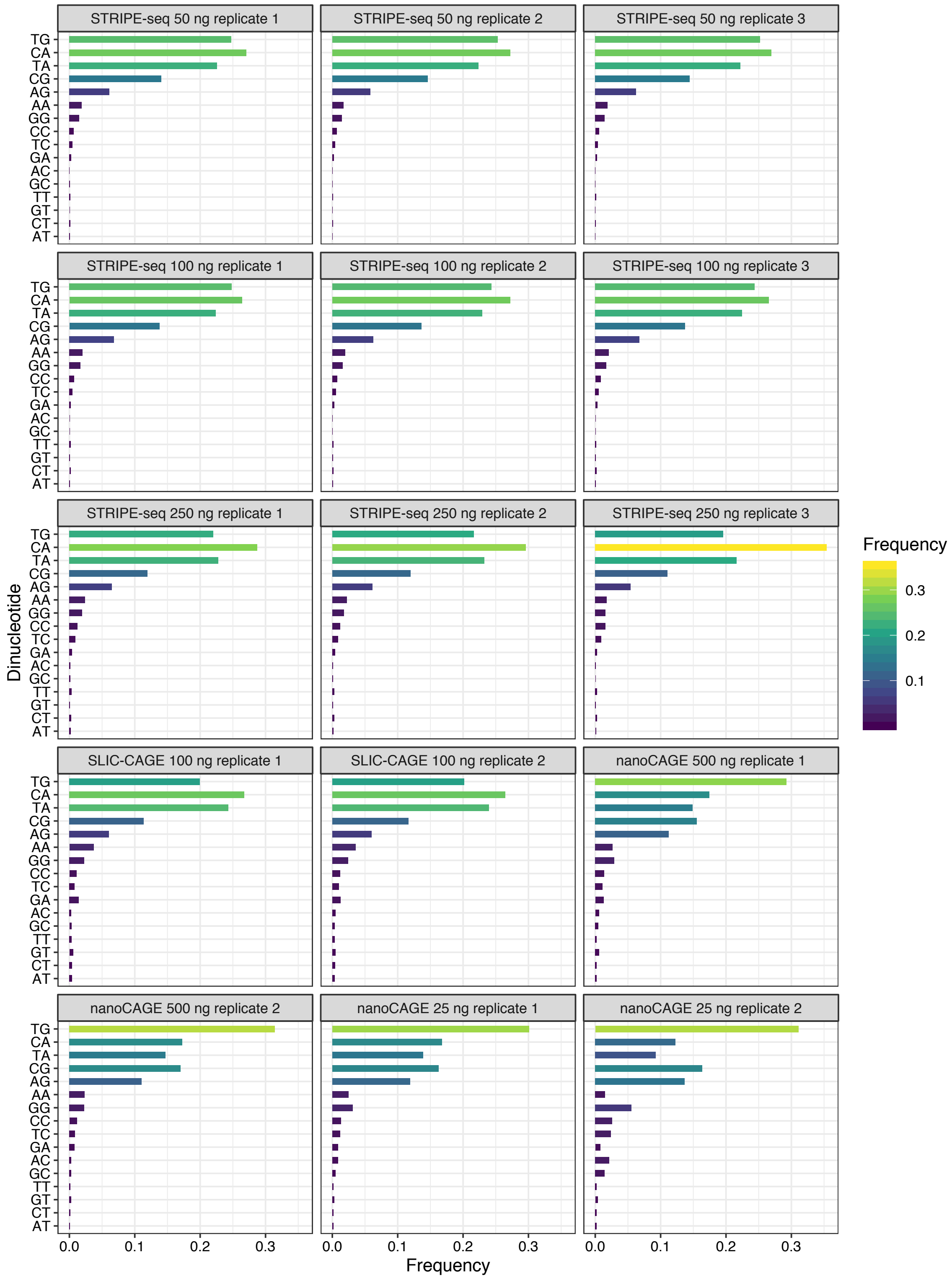


**Supplemental Figure 14. Dinucleotide frequencies at yeast TSSs identified by STRIPE-seq, SLIC-CAGE, and nanoCAGE**

Dinucleotides are in descending order based on their average frequency across all samples.

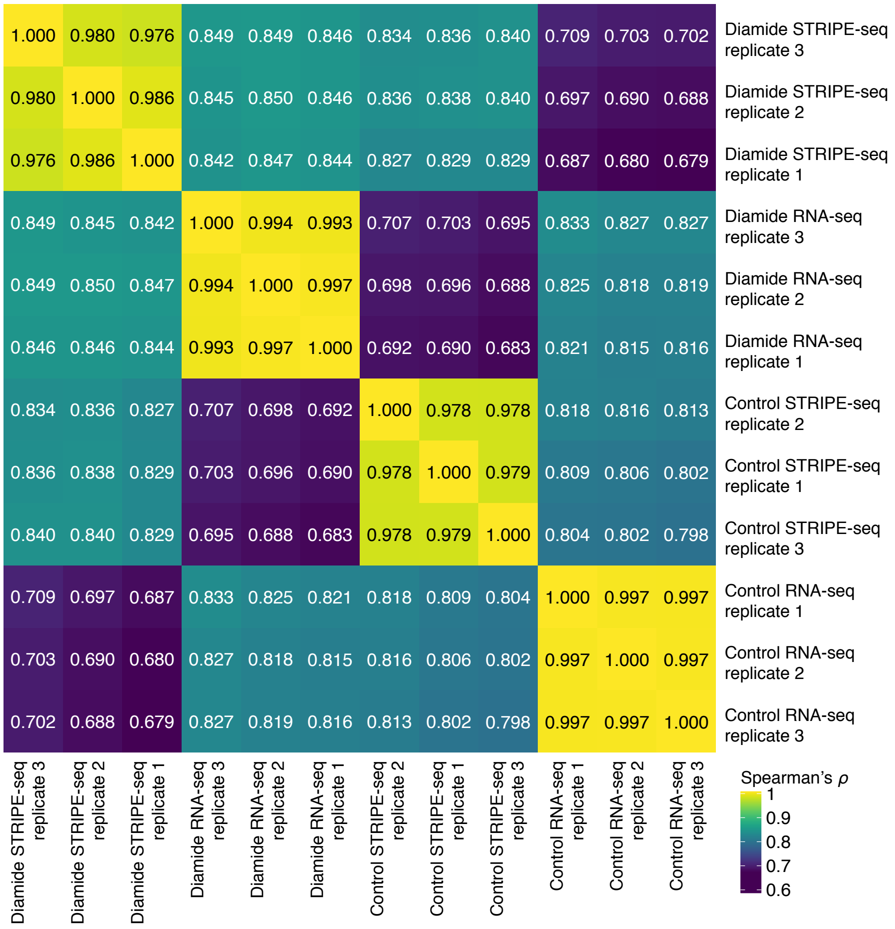


**Supplemental Figure 15. Correlation of yeast control and diamide STRIPE-seq and RNA-seq signal**

STRIPE-seq and RNA-seq fragments within transcripts were counted, compared by Spearman correlation analysis, and plotted as a hierarchically clustered heatmap.

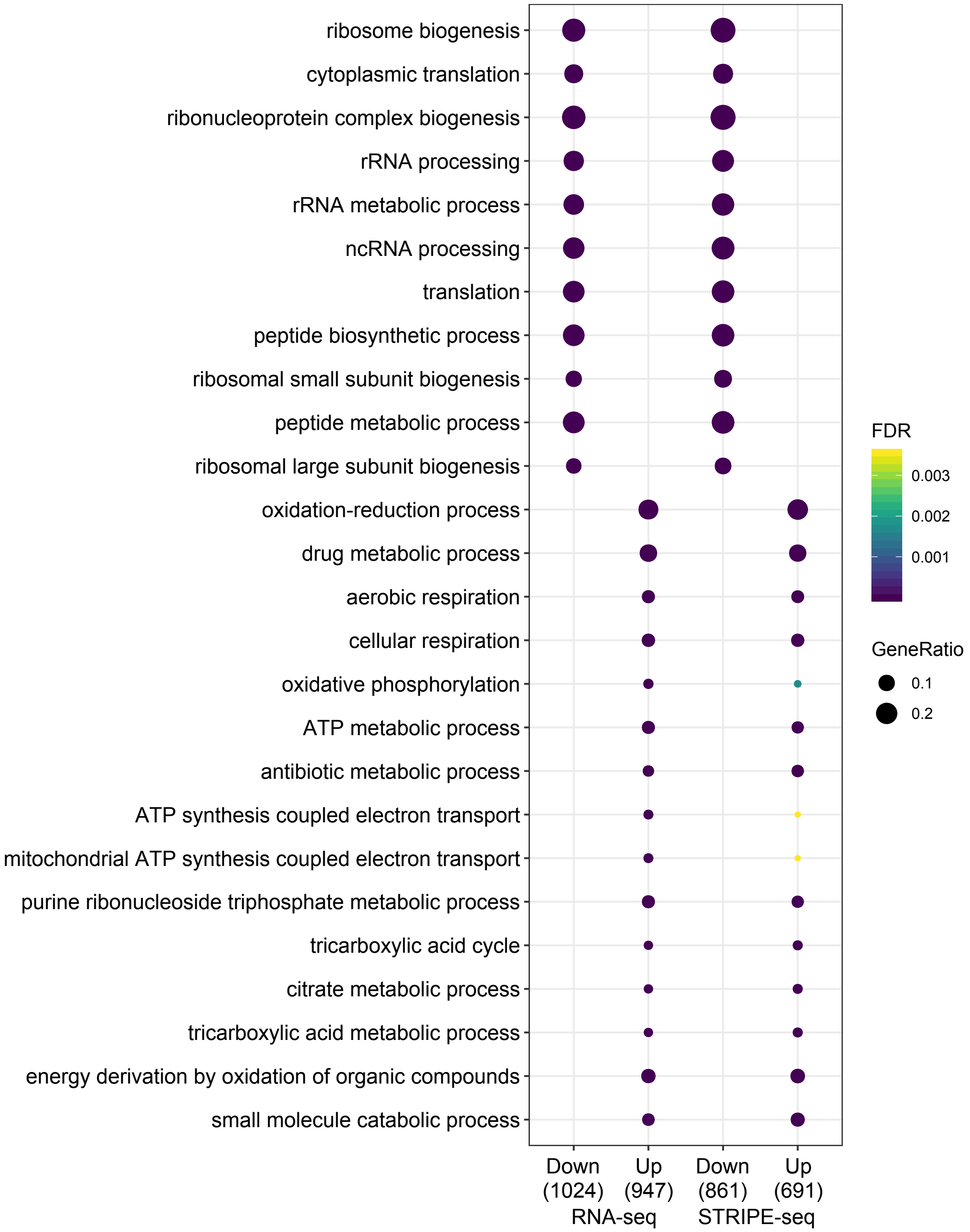


**Supplemental Figure 16. GO biological process analysis of DEGs in STRIPE-seq and RNA-seq data for control and diamide-treated yeast**

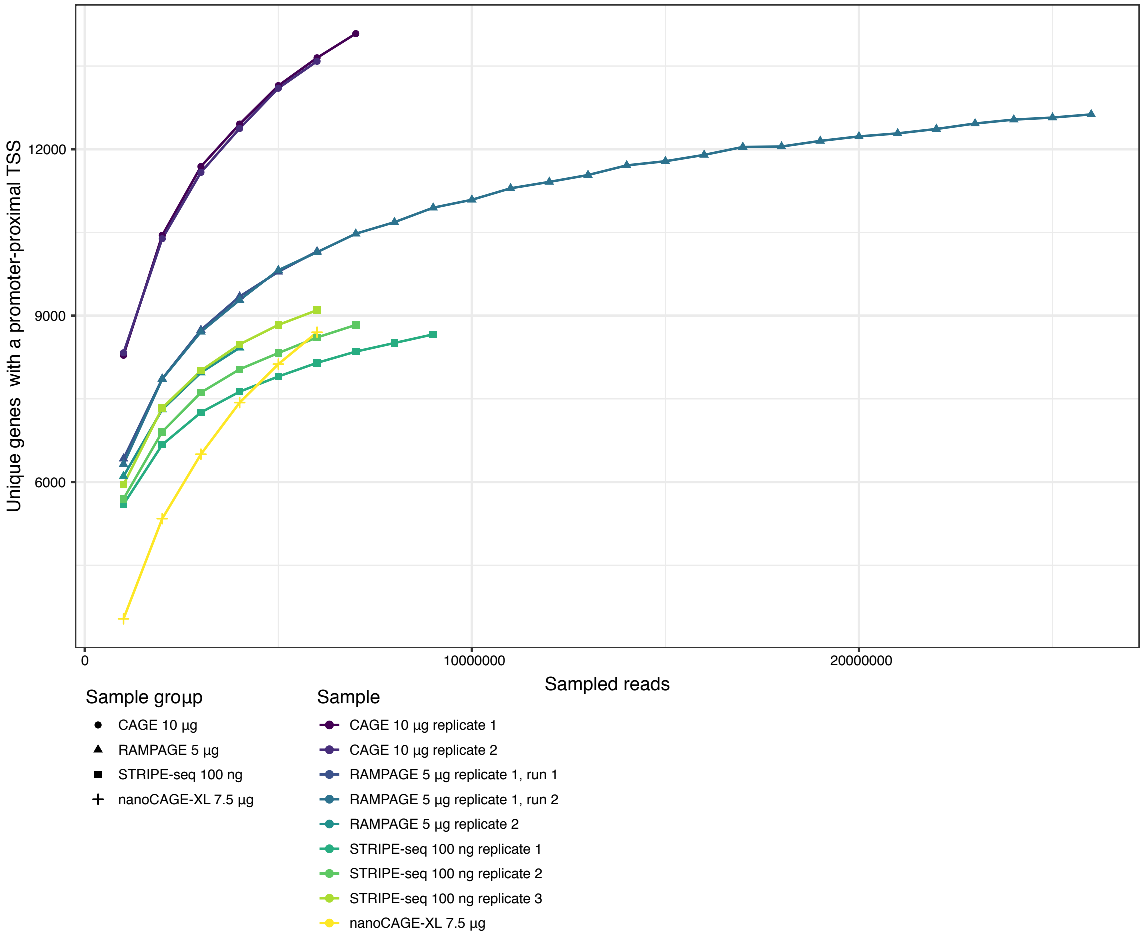


**Supplemental Figure 17. Saturation analysis of K562 TSS mapping datasets**

Analysis of sequencing saturation in STRIPE-seq, RAMPAGE, CAGE, and nanoCAGE-XL datasets. rRNA was computationally removed from FASTQ files and reads were aligned to the human GRCh38.p13 genome assembly. The given numbers of mappable reads were sampled from the BAMs, PCR duplicates were removed, and only primary alignments with properly paired mates were retained. TSSs were then called using TSRchitect and annotated using ChIPseeker.

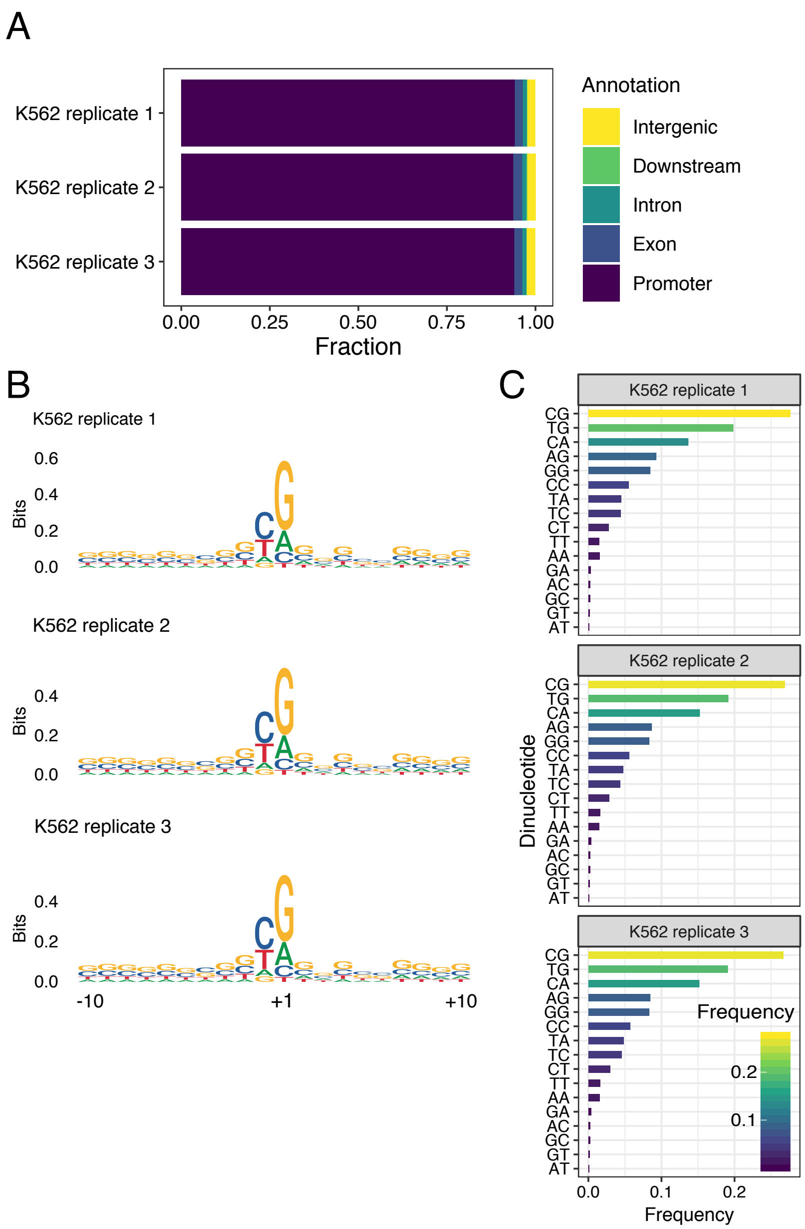


**Supplemental Figure 18. Genomic context analysis of K562 TSSs detected by STRIPE-seq**

(A) Genomic distribution of K562 TSSs identified by STRIPE-seq. (B) Sequence logos for K562 TSSs identified by STRIPE-seq. (C) Dinucleotide frequencies at K562 TSSs identified by STRIPE-seq. Dinucleotides are in descending order based on their average frequency across all samples.

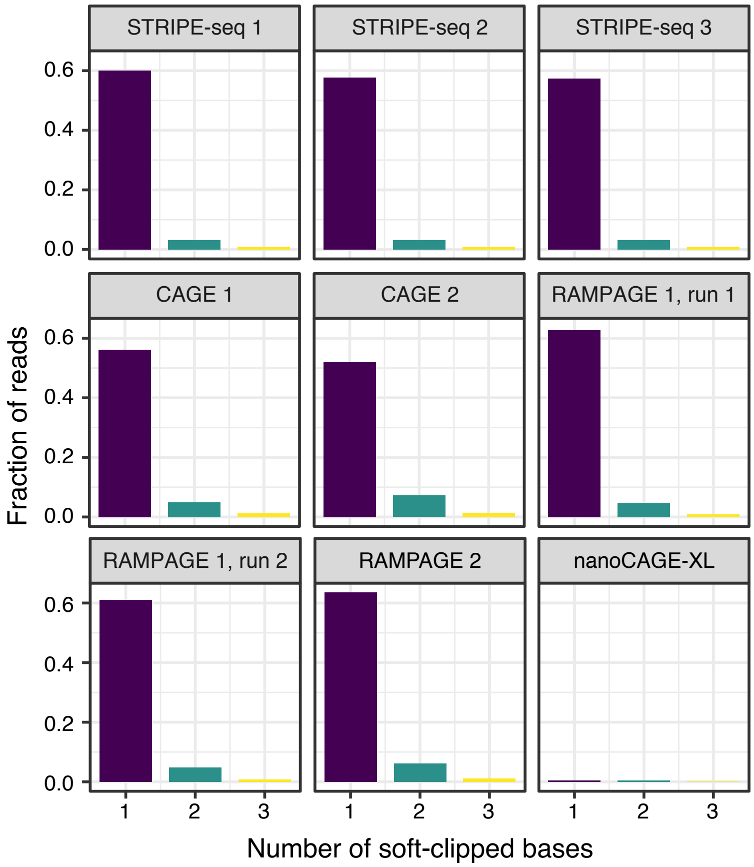


**Supplemental Figure 19. Analysis of soft-clipped base prevalence in human TSS mapping datasets**

Bar charts showing the frequencies of STRIPE-seq, CAGE, RAMPAGE, and nanoCAGE-XL R1 reads with the given number of soft-clipped bases extending upstream from the called TSS. Reads with more than 3 soft clipped-bases were infrequent in all methods and were thus not analyzed further.

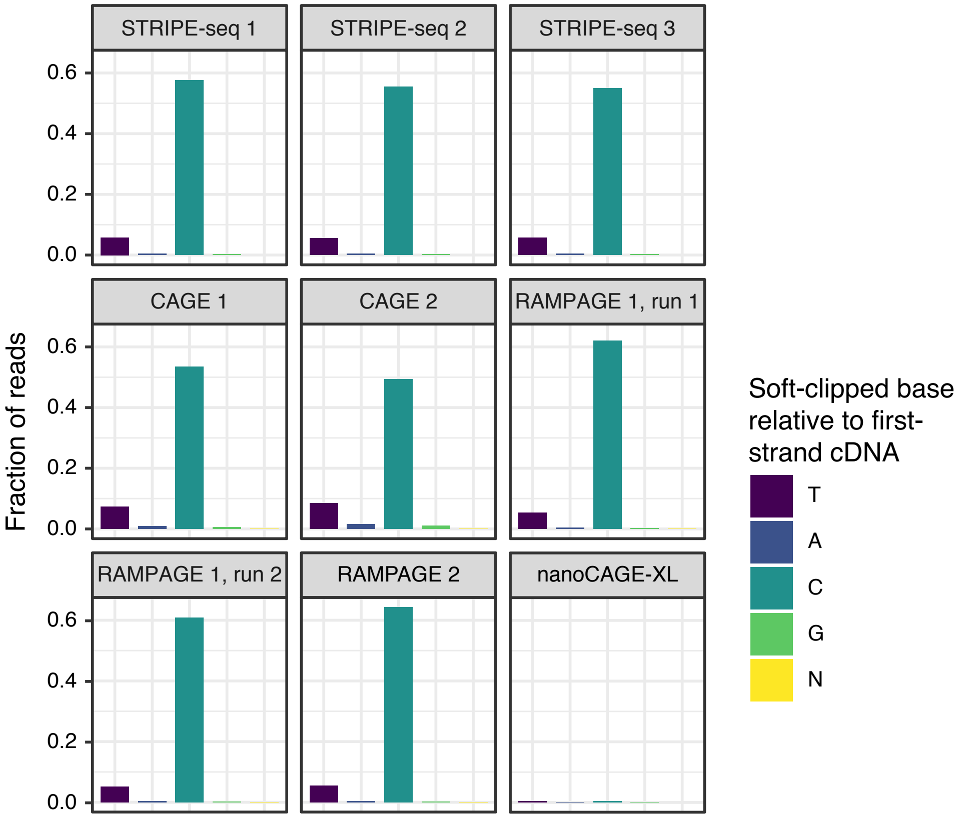


**Supplemental Figure 20. Sequence distribution of soft-clipped bases in human TSS mapping datasets**

Bar charts showing the frequencies of soft-clipped bases in STRIPE-seq, CAGE, RAMPAGE, and nanoCAGE-XL at the position directly upstream of the called TSSs. Bases displayed are relative to the first-strand cDNA.

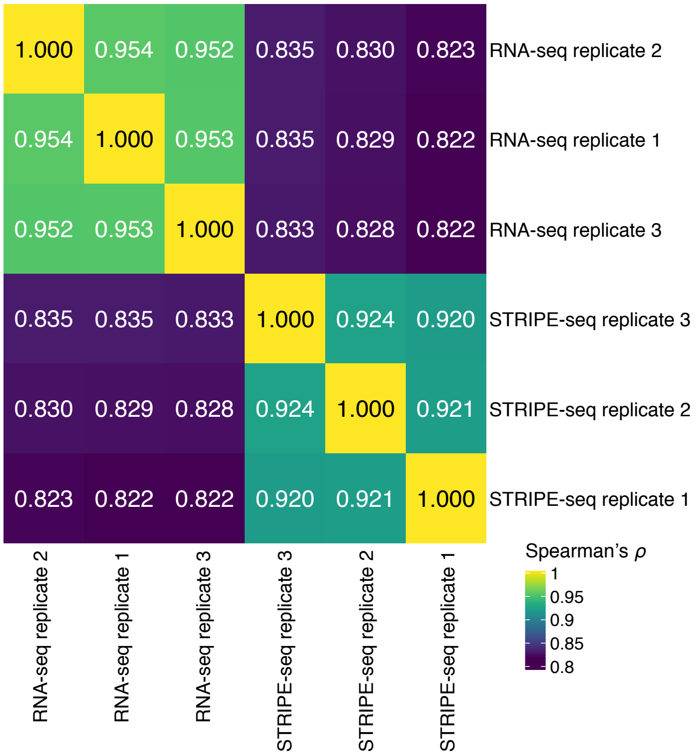


**Supplemental Figure 21. Comparison of K562 STRIPE-seq and RNA-seq transcript abundance measurements**

STRIPE-seq and RNA-seq fragments within transcripts were counted, compared by Spearman correlation analysis, and plotted as a hierarchically clustered heatmap.

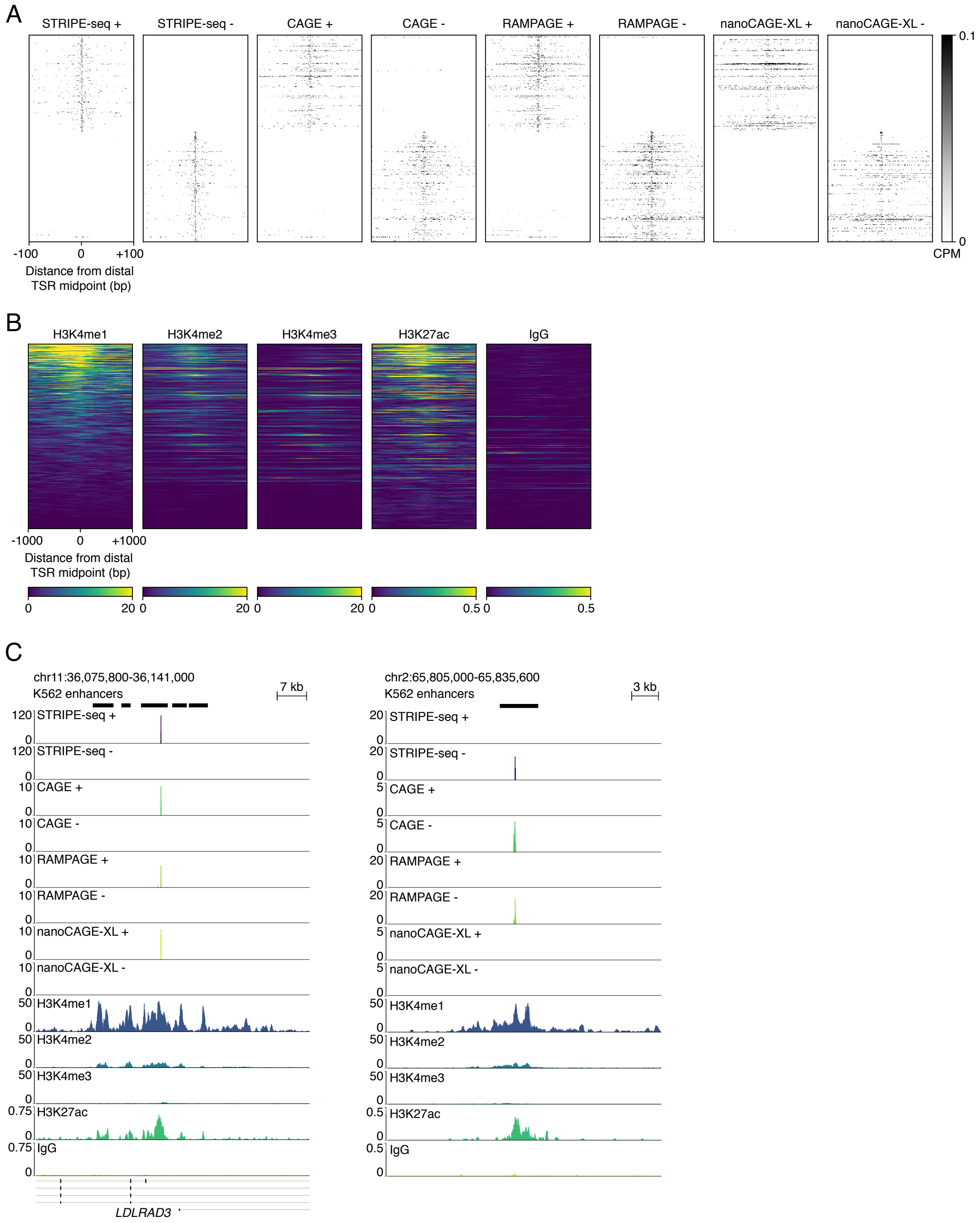


**Supplemental Figure 22. Analysis of K562 distal TSRs**

(A) Heatmaps of CPM-normalized STRIPE-seq, CAGE, RAMPAGE, and nanoCAGE-XL signal at 513 distal (≥ 1 kb from an annotated TSS) TSRs detected in all three K562 STRIPE-seq replicates. Replicate 1 is shown for all data types. (B) Heatmaps of spike-in-normalized H3K4me1/2/3, H3K27ac, and IgG control CUT&Tag signal at distal K562

TSRs. Heatmaps are sorted descending by average H3K4me1 signal. (C) Genome browser-style tracks showing CPM-normalized replicate 1 STRIPE-seq, CAGE, RAMPAGE, and nanoCAGE-XL signal, spike-in-normalized H3K4me1/2/3, H3K27ac, and IgG control CUT&Tag signal, and EnhancerAtlas 2.0 K562 enhancer annotations at two regions of the human genome.

| | Paper | | RNA Input (ng) | | Steps | | Time (hours) | | | Costs (USD) | | | | Commerical Kits | |
|---|---|---|---|---|---|---|---|---|---|---|---|---|---|---|---|
| | PMID | Year | Minimum | Maximum | Required | Optional | Required | With Optional | Overnight Incubations | Startup | Required Per Sample | With Optional | Additional After Pooling | Website | Per Sample Cost |
| **RAMPAGE** | 24510412 | 2013 | 5000 | 5000 | 12 | - | 13.5 | - | 15 | 5844 | 55.4 | - | 48.79 | - | - |
| **nAnT-iCAGE** | 24927836 | 2014 | 5000 | 5000 | 22 | 1 | 18.75 | 21.75 | 16 | 9071 | 105.1 | 119.81 | - | https://cage-seq.com/cage_kit/ | 225 |
| **nanoCAGE** | 28349422 | 2017 | 50 | 500 | 9 | 3 | 12.5 | 15 | - | 3732 | 18.69 | 29.28 | 37.82 | - | - |
| **SLIC-CAGE** | 31305534 | 2019 | 1 | 100 | 33 | - | 36 | - | 16 | 7340 | 113.84 | - | - | - | - |
| **STRIPE-seq** | - | 2020 | 50 | 200 | 6 | - | 5 | - | - | 2294 | 11.73 | - | - | - | - |

### Supplemental Figure 23. Cost and time comparison of TSS mapping methods

A summary of the time and cost analysis of TSS mapping methods presented in Supplemental Table 4.
